# Steering cell-state and phenotype transitions by causal disentanglement learning

**DOI:** 10.1101/2024.08.16.607277

**Authors:** Chengming Zhang, Zexi Chen, Yuanxiang Miao, Yun Xue, Deyu Cai, Weifeng Guo, Hongbin Ji, Kazuyuki Aihara, Luonan Chen

**Affiliations:** International Research Center for Neurointelligence, The University of Tokyo Institutes for Advanced Study, The University of Tokyo, Tokyo 113-0033, Japan; Key Laboratory of Systems Biology, Shanghai Institute of Biochemistry and Cell Biology, Center for Excellence in Molecular Cell Science, Chinese Academy of Sciences, Shanghai 200031, China; School of Life Science and Technology, ShanghaiTech University, Shanghai 201210, China; School of Electrical and Information Engineering, Zhengzhou University, Zhengzhou 450001, State; Key Laboratory of Intelligent Agricultural Power Equipment, China and State Key Laboratory of Oncology in South China; Key Laboratory of Systems Health Science of Zhejiang Province, School of Life Science, Hangzhou Institute for Advanced Study, University of Chinese Academy of Sciences, Chinese Academy of Sciences, Hangzhou 310024, China; Guangdong Institute of Intelligence Science and Technology, Hengqin, Zhuhai, Guangdong 519031, China

**Keywords:** Causal disentanglement, Information flow, State transition, System control, Interpretive framework, Deep learning

## Abstract

Understanding and manipulating cell-state and phenotype transitions is essential for advancing biological research and therapeutic treatment. We introduce CauFinder, an advanced framework designed to accurately identify causal regulators of these transitions and further precisely steer such transitions by integrating causal disentanglement modelling with network control based solely on observed data. By leveraging do-calculus and optimizing information flow metrics, CauFinder can distinguish causal factors from spurious ones, ensuring precise control over desired state transitions. One significant advantage of CauFinder is its ability to identify those variables causally affecting the cell-state/phenotype transitions among all observed variables, both theoretically and computationally, leading to the identification of their master regulators when combined with network control. Consequently, by employing a counterfactual algorithm, CauFinder is able effectively to facilitate desirable state transitions or steer these transitional trajectories/paths by modulating these causal drivers. Beyond its theoretical advantages, CauFinder outperforms existing approaches computationally in both simulated and real-world settings. CauFinder is able to not only reveal natural biological transition processes such as (a) cell differentiation, (b) lung adenocarcinoma (LUAD) to lung squamous cell carcinoma (LUSC) transdifferentiation and (c) drug-sensitive to drug-resistant transitions but also identify the causal regulators of their reverse transition processes, such as (A) cell dedifferentiation, (B) LUSC to LUAD transdifferentiation and (C) drug-resistant to drug-sensitive transitions. These findings highlight its superior ability to causally uncover essential regulatory mechanisms and accurately steer cell-state/phenotype transitions, thus providing novel therapeutic strategies.

## Introduction

Transitions in cell states and phenotypes, such as changes in cell types during development or shifts from healthy to diseased tissues, are fundamental processes in biological systems and crucial for understanding biology and diseases. Clarifying transition processes and identifying causal drivers behind these transitions is vital for uncovering the mechanisms of biological evolution, health, and adaptability. However, fully elucidating and manipulating these transitions remains a challenge. Identifying the causal factors that drive these transitions is particularly crucial, as they play a critical role in the stability and transformation of biological systems^1^. Therefore, precisely identifying these causal control points/drivers and understanding their functions across various biological processes and under different environmental conditions are essential for advancing our comprehension of biological systems and enabling precise interventions^2^. This understanding holds substantial potential for applications in cell fate determination, tissue regeneration, and disease treatment, demonstrating broad implications for biomedicine.

To control a biological transition by identifying its key drivers from the observed data, various traditional methods have been developed. Machine learning models, such as logistic regression and random forests, assess feature importance during model fitting, but primarily focus on correlations rather than causative mechanisms, thus potentially leading to misidentification of true drivers^3^. Similarly, statistical models, including differential expression analyses like t-tests and non-parametric tests, identify significant disparities but do not establish causality. Network-based approaches and advancements in deep learning offer new insights into biological transitions. For example, WMDS.net^4^ identifies key regulators in transcriptional networks by integrating node degree and gene co-expression differences to find a minimum dominating set (MDS)^5^ of driver nodes. Meanwhile, the MOVE^6^ framework employs multi-omics variational autoencoders to discover drug-omics associations in type 2 diabetes, highlighting complex interactions like those between metformin and the gut microbiota. Similarly, CEFCON^7^ uses graph neural networks with attention mechanisms to infer cell lineage-specific gene regulatory networks from single-cell RNA data, applying control theory to identify critical regulators of cell fate. Despite their advances, these methods primarily focus on association rather than causation, and while some leverage control theory to identify key regulators, they often do not fully elucidate the causal mechanisms underlying biological processes. This fact highlights the need for data-driven approaches that can integrate both causality and control theory so as to provide effective interventions.

The profound significance of causality in deciphering and manipulating cell-state and phenotype transitions has been widely acknowledged, offering a deeper and more transparent understanding of biological systems and facilitating the development of interventions^8–10^. While association analyses can yield preliminary insights, they may inadvertently lead to inaccurate conclusions due to the potential interference of confounding factors^11–13^. Although Randomized Controlled Trials (RCTs) have been historically esteemed as the gold standard for causal inference, they have limitations, such as constrained applicability, potential ethical issues, and substantial demands on time and resources^14–17^. Consequently, our focus shifts to computational techniques, which utilize do-calculus to simulate interventions. These techniques align with the objectives of RCTs but without inheriting their limitations, thus enabling a more adept identification of authentic relationships^18,19^. The concept of ’Causal Emergence’ reveals that the collective impact of individual features is significantly amplified, highlighting the need to integrate these collective effects into our analyses^20,21^. However, many existing causality analysis methods grounded in Directed Acyclic Graphs (DAGs) focus on relations between individual variables and outcomes^22^, neglecting the collective effect of groups of variables on phenotypes and also the feedback (loop) effect.

To address these challenges, we developed herewith CauFinder, a advanced deep learning-based causal model designed to identify a subset of master regulators that collectively exert a significant causal impact during cell-state or phenotype transitions from the observed data. CauFinder elucidates state transitions by identifying causal factors within a latent space and quantifying causal information flow from latent features to state predictions. It can theoretically identify and circumvent confounders using the backdoor adjustment formula^23^. Beyond do-calculus intervention, CauFinder employs SHapley Additive exPlanations (SHAP)^24^ values and gradient calculations to identify key causal features in the original data space and quantify their causal strength and direction on state transitions. Moreover, CauFinder can identify the master regulators that control these transitions, by integrating causal strength with the feedback vertex set (FVS)^25,26^ method on a gene regulatory network constructed from the observed data. By employing a counterfactual algorithm that balances the proximity to the desired outcome and minimal perturbation to the causal variables, CauFinder effectively modulates these features to achieve controlled and desirable state transitions. Our results demonstrate that CauFinder outperforms traditional methods in all the simulated, single-cell, and bulk benchmark datasets in identifying causal drivers. Furthermore, our analyses found that CauFinder is able to not only reveal natural biological transition processes such as (a) cell differentiation and (b) lung adenocarcinoma (LUAD) to lung squamous cell carcinoma (LUSC) transdifferentiation and (c) drug-sensitive to drug-resistant transitions but also identify the causal regulators of their reverse transition processes, such as (A) cell dedifferentiation, (B) LUSC to LUAD transdifferentiation and (C) drug-resistant to drug-sensitive transitions. These findings highlight CauFinder’s robustness and versatility in uncovering causal regulatory mechanisms, paving the way for effective interventions in precision medicine and biotechnology.

## Results

### Overview of CauFinder

The primary goal of CauFinder is to dissect and manipulate biological processes by precisely identifying and controlling master regulators essential for cell state or phenotype transitions. CauFinder comprises two key components (Fig. 1a). Initially, the framework utilizes a causal decoupling/disentangling model within a variational autoencoder (VAE) architecture to distill complex datasets into distinct causal and spurious elements. This model isolates features that directly influence cell-state phenotype transitions (Fig. 1b and 1c), distinguishing genuine causal relations from mere correlations: a critical step for accurate biological interpretation and targeted interventions. Notably, CauFinder has the ability to theoretically and computationally identifying the partial variables causally affecting the cell-state or phenotypes among all observed variables.

**Fig 1:**
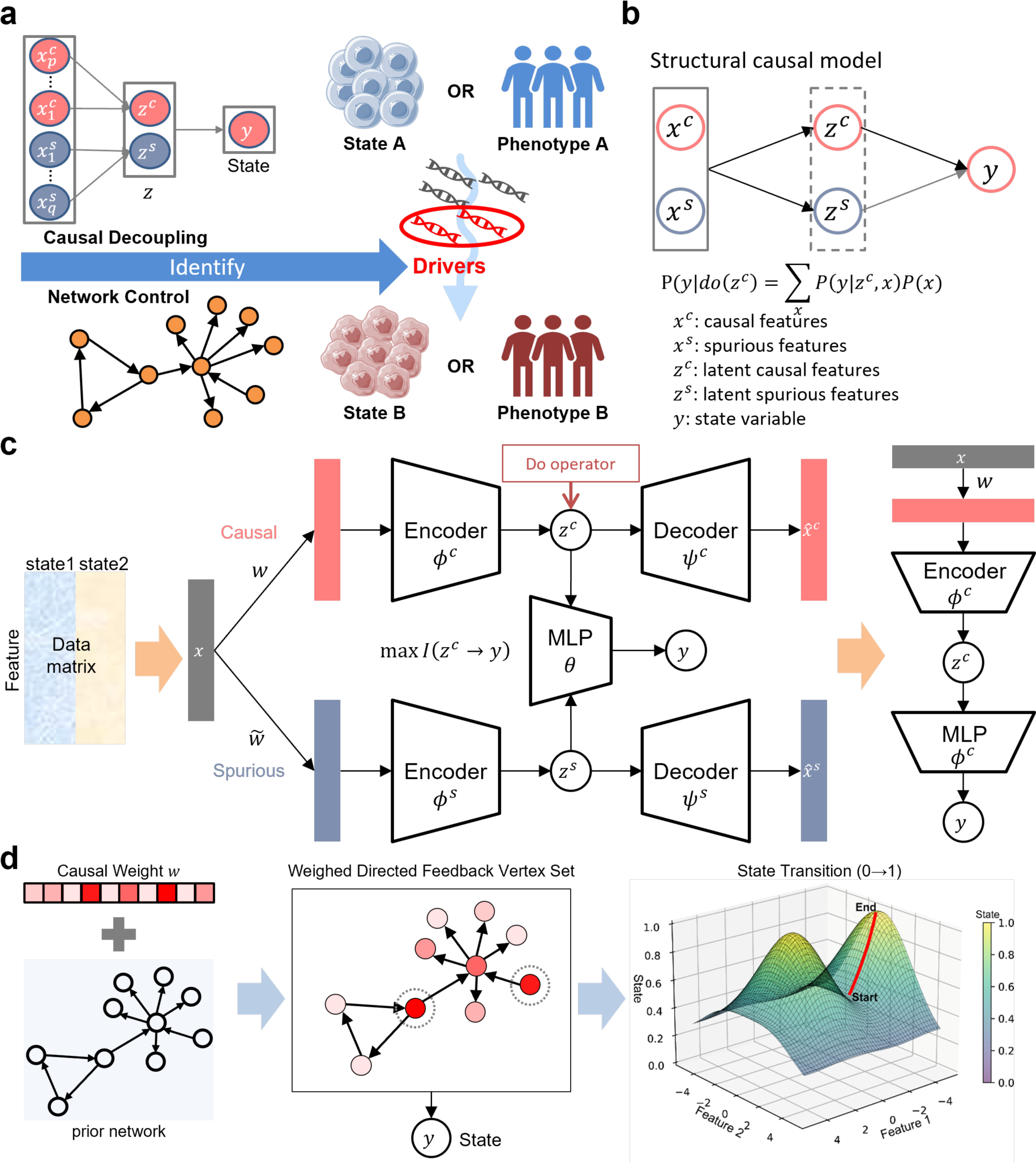
Overview of the CauFinder framework. **a**, Schematic representation of the CauFinder framework for steering state or phenotype transitions by identifying key drivers through causal disentanglement and network control. **b**, Structural causal model showing the decomposition of original features *x* = {*x*^*c*^, *x*^*s*^} into their latent representations *z* = {*z*^*c*^, *z*^*s*^}, with *y* denoting the state or phenotype. **c**, Causal decoupling model constructed on a variational autoencoder (VAE) framework to identify causal drivers influencing state or phenotype transitions. Features *x* are processed through a feature selection layer, encoded to create a latent space *z*, segmented into causal (*z*^*c*^) and spurious (*z*^*s*^) components. This latent space is decoded to reconstruct the original features *x* as *x̂* and to predict the phenotype *y*. The model strives to maximize the causal information flow, *I*(*z*^*c*^ → *y*), from *z*^*c*^ to *y*, thus delineating the causal pathways from *x* to *y* via *z*^*c*^ and identifying the causal drivers for precise transition control. **d**, Master regulator identification via causality-weighted features and network control. Techniques including SHAP and gradient are used to assign causality weights to features within the causal path defined in **c**, aiding in the isolation of causal features for integration with prior network insights. Weighed directed feedback vertex set is then employed to pinpoint master regulators critical for directing state or phenotype transitions through counterfactual generation for causal state transition, thereby establishing the foundation for targeted interventions.

Following causal analysis, CauFinder integrates the delineated causal features with prior network knowledge using a novel weighted feedback vertex set (FVS) method. This approach systematically assesses the impact of potential master regulators within the network, pinpointing those with substantial influence on desired transitions (Fig. 1d). By leveraging this technique, CauFinder is able to identify and rank master regulators based on their control efficacy, equipping researchers with a potent tool for targeted genetic or pharmacological interventions.

This dual-component strategy enables CauFinder to elucidate underlying mechanisms driving phenotype changes and to orchestrate precise and effective biological interventions. The integration of causal discovery with network theory provides a robust framework for advancing our understanding of complex biological systems and developing targeted therapies that can significantly modify disease states or enhance desirable phenotypic traits. The innovative merger of causal inference and network control in CauFinder represents a significant advancement in our ability to understand and directly manipulate cellular states at a molecular level, paving the way for next-generation therapies that are both more effective and precise.

### Benchmark evaluation of CauFinder on simulated datasets

Evaluating methods for discovering causality in data is inherently challenging due to the absence of ground truth. To address this issue, we first constructed simulated datasets, which served as a basis for assessing the efficacy of CauFinder and other competing methods in identifying causal relations. The design of the simulated data is illustrated in Fig. 2a and detailed in Supplementary Note 1. The dataset includes observed variables with direct causal effects (*x*^*c*^), spurious correlations (*x*^*s*^), and the outcome variable (*y*). The unobserved variable *u* acts as a confounder, influencing both *x*^*s*^and *y*. The primitive causal variable *c* directly affects *x*^*c*^. In generating the dependent variable *y*, we adjusted the parameters *λ* and *ε* to control the causal strength and noise scale, creating different scenarios. Our objective is to use this simulated dataset to compare the ability of CauFinder and other competing methods to identify *x*^*c*^ among all the observed variables.

**Fig 2:**
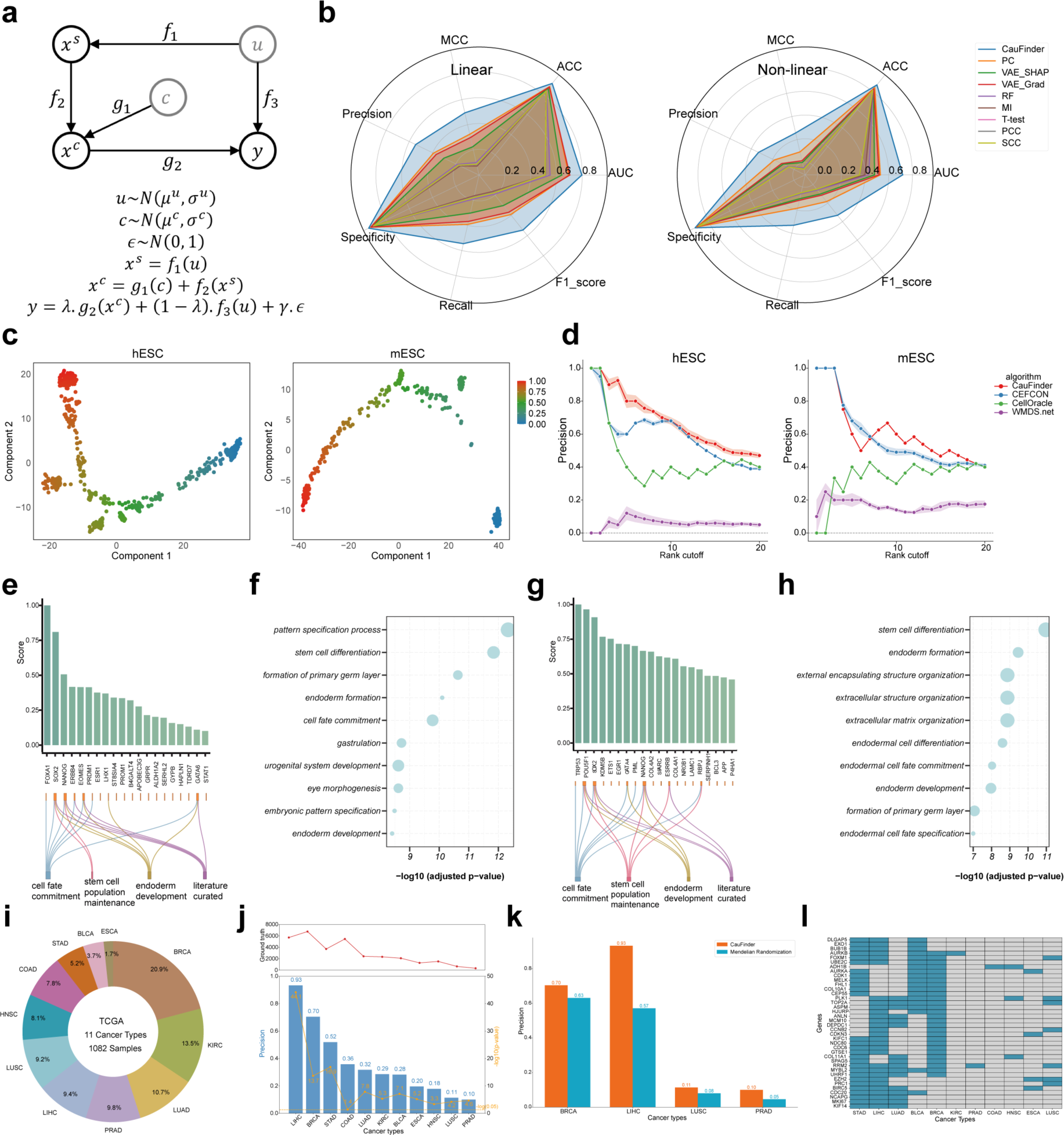
Evaluation of method performance in simulated and real-world datasets. **a**, Construction of simulation data. Here, *x*^*s*^ represents the observed spurious variables, *x*^*c*^denotes the observed causal variables, and *y* is the outcome variable influenced by both *x*^*c*^ and unobserved variables *u*. **b,** Performance evaluation of CauFinder and comparative methods on simulated data in both linear (left) and nonlinear (right) scenarios. Radar charts show the average values of evaluation metrics for each method under different noise levels and causal strengths. **c,** Pseudotime trajectories for hESC and mESC. The plots display the differentiation trajectories of human embryonic stem cells (hESC) and mouse embryonic stem cells (mESC) based on pseudotime analysis performed using Monocle 3. The color gradient represents pseudotime, with red indicating early stages and blue indicating later stages of differentiation. Component 1 and Component 2 are the first two dimensions obtained from UMAP, used to visualize the cell differentiation trajectories. **d,** Performance on the hESC and mESC datasets for each method, evaluated in terms of the precision of the top-*k* predicted genes among all known genes in the ground-truth gene set. The methods compared include CauFinder, CEFCON, CellOracle, and WMDS.net. All the results with *k* ranking from 1 to 20 were reported. The shaded area represents the variation (mean ± s.d.) of precision over 10 repeats. **e, g,** The top 20 predicted driver regulators in the hESC (**e**) and mESC (**g**) are ranked in descending order based on their normalized scores from CauFinder. The genes corresponding to each ground-truth gene set are displayed below the bar chart. **f, h,** The top enriched GO terms for all genes predicted as driver regulators in hESC (**f**) and mESC (**h**). The size of the dots represents the number of driver genes enriched in each term, and the terms are ranked by −log10(P value). P values were adjusted using the Benjamini–Hochberg method in the clusterProfiler package. **i,** Donut chart showing the distribution of paired samples from different cancer types in the TCGA dataset analyzed. The size of each sector represents the proportion of samples for each cancer type. **j**, Top line chart representing the total number of pathogenic genes documented in the literature for each cancer type, as sourced from the DisGeNET database. The bottom bars and line chart showing the precision (blue bars) of pathogenic genes identified by CauFinder in each cancer type, compared against literature-documented pathogenic genes, and the - log10(p-value) (orange line). The horizontal axis denotes different cancer types, the left vertical axis shows precision, and the right vertical axis shows the -log10(p-value). **k,** Bar chart comparing the precision of pathogenic gene identification by CauFinder and Mendelian Randomization (MR) in four cancer types: breast invasive carcinoma (BRCA), liver hepatocellular carcinoma (LIHC), lung squamous cell carcinoma (LUSC), and prostate adenocarcinoma (PRAD). Precision is calculated by comparing the genes identified by each method with the pathogenic genes documented in the literature. The orange bars represent the precision of CauFinder, and the blue bars represent the precision of Mendelian Randomization. **l,** Heatmap illustrating the occurrence of genes identified by CauFinder as drivers in at least three different cancer types. The heatmap includes the top genes sorted by their frequency across 11 cancer types. The horizontal axis represents the cancer types, and the vertical axis lists the genes. Blue indicates the presence of the gene in a particular cancer type, while grey indicates its absence.

The competing methods employed for this comparison include the Peter and Clark (PC) algorithm^27^, random forest (RF), T-test, variational autoencoder (VAE), mutual information (MI), Pearson correlation coefficients (PCC) and Spearman correlation coefficients (SCC) (Supplementary Note 2). For the simulated data, we evaluated the performance of each method under different combinations of noise levels and causal strengths in both linear and nonlinear scenarios. The evaluation metrics used for simulated datasets are detailed in Supplementary Note 3. The choice of the number of causal drivers was guided by the strategies detailed in Supplementary Note 4. However, CauFinder consistently outperforms the other methods, regardless of whether the data are linear or nonlinear. Notably, the PC algorithm also performed well across all metrics, securing the second place, which underscores the superiority of causal methods over correlation-based methods in identifying causal features. Specifically, we presented the AUC and F1-scores of each method under different noise levels and causal strengths in both linear and nonlinear scenarios (see Supplementary Figs. 1-4). We observed that as noise increased or causal strength decreased, the performance of all methods declined, which is expected. Nevertheless, CauFinder consistently outperformed other methods in each specific combination.

Furthermore, we analyzed CauFinder’s performance in causal decoupling on the simulation dataset. In the model’s latent space, the information flow of the causal dimension was significantly higher than that of the spurious dimension (Supplementary Fig. 5a, b). Compared to spurious features, the model assigned higher weights to causal features (Supplementary Fig. 5c). The original space could not achieve a clear differentiation between classes (Supplementary Fig. 5d), while the sample distribution based on the causal dimension in the latent space indicated a clearer differentiation between classes compared to the spurious latent space (Supplementary Fig. 5e). These results demonstrated that CauFinder can accurately disentangle causal and spurious factors in both the original and latent space.

We also explored the impact of the dimension of causal latent spaces on model performance (Supplementary Fig. 6). The results show that while the variation is not significant, the performance is optimal when the dimension of causal latent spaces is set to 2 (out of a total of 10 dimensions). To verify the critical role of causal information flow components in identifying causal features, we conducted ablation experiments on simulated data (Supplementary Figs. 7-8). The results indicated that the model with causal information flow (CauFinder) outperformed the model without causal information flow (CauFinderNC), further highlighting the importance of causal information flow in identifying causal features.

### Benchmark evaluation of CauFinder on single-cell and bulk datasets

To further validate the performance of CauFinder, we evaluated it on real-world datasets, specifically using data from embryonic stem cells (ESC) for single-cell data and The Cancer Genome Atlas (TCGA) for bulk data.

We first evaluated the performance of CauFinder in identifying driver regulatory genes using mouse embryonic stem cell (mESC) and human embryonic stem cell (hESC) datasets. We defined a comprehensive gold standard gene set from CEFCON^7^ associated with cellular fate and ESC development to benchmark our results. These gene groups included ’cell fate commitment’, ’stem cell population maintenance’, ’endoderm development’ and ’literature curated’ sets, ensuring a comprehensive basis for evaluation. Figure 2c visualizes the pseudotime trajectories for hESC and mESC datasets, with the color gradient representing the progression of pseudotime from early to late stages of differentiation, generated using Monocle^28^. We used the top 25% and bottom 25% of cells based on pseudotime as two distinct states for input labels in CauFinder’s analysis. We quantified accuracy by the precision of the top-*k* predictions and compared CauFinder with several benchmark methods, including CEFCON, CellOracle^29^, and WMDS.net (see Supplementary Note 2). In the hESC dataset, CauFinder demonstrated superior precision, notably at early rank cutoffs, highlighting its ability to swiftly and accurately identify essential regulatory genes. CEFCON performed commendably, especially at mid-rank cutoffs, but did not reach the early precision levels of CauFinder. In contrast, CellOracle and WMDS.net were markedly less precise, with WMDS.net showing the lowest precision across all methods (Fig. 2d). In the mESC dataset, CauFinder matched CEFCON at initial rank cutoffs and gradually surpassed it, indicating superior precision across a wider array of gene identifications. CellOracle improved at higher rank cutoffs, while WMDS.net consistently underperformed, underscoring CauFinder’s enhanced capabilities in complex genomic analysis (Fig. 2d). These results underscore CauFinder’s substantial advantages in identifying driver regulatory factors, providing a solid foundation for further biological research.

We next examined the top 20 driver regulators identified by CauFinder in hESC and mESC datasets, noting that nearly half of them aligned with established ground-truth gene sets (Fig. 2e, g). Specifically, the transcriptional factors (TFs) SOX2 (rank 2/20 in hESC and 3/20 in mESC) and NANOG (rank 3/20 in hESC and 9/20 in mESC) are recognized pluripotency factors^30^ and have been reported to form a cross-regulatory feedforward loop in both species^31^. Additional well-known pluripotency factors, including EOMES and GATA6 (rank 5/20, 19/20) for hESC, as well as POU5F1 and GATA4 (rank 2/20, 7/20) for mESC, also received high scores. Among these, GATA4 and GATA6 have been previously shown to induce selective differentiation to primitive endoderm^32^. Beyond these TFs, CauFinder also identified key non-TF driver regulators such as ALDH1A2, PROM1, PML, and KDM5B, which play crucial roles in ESC differentiation. For instance, PML belongs to a transcriptional repressive complex with OCT4 and NANOG and is essential for the self-renewal and pluripotency of ESCs^33^. ALDH1A2 is involved in retinoic acid (RA) synthesis and signalling, essential for cell differentiation^34^. PROM1 is a marker for various stem and progenitor cells^35–37^ and is a key regulator ensuring proper responses to extracellular signals in stem cells^38^. KDM5B belongs to the KDM5 subfamily of histone lysine-specific demethylases^39^. It inhibits the expression of pluripotency genes, SOX2 and NANOG^40^, and has been identified as a major downstream target of the transcription factor NANOG, which plays a central role in the initiation and maintenance of ESC pluripotency and self-renewal^41^. We then performed biological pathway enrichment analyses of the driver regulators identified by CauFinder (Gene Ontology: biological process). We selected the top 10 pathways and further ranked them based on their log-transformed P-values (Fig. 2f, h). Several expected pathways shared between samples were observed: stem cell differentiation and endoderm formation were functionally enriched by driver regulators from both mESC and hESC. Pattern specification processes were driven by driver regulators from hESC (Fig. 2f), while extracellular structure organization was driven by driver regulators from mESC (Fig. 2h).

CauFinder demonstrates the capability to identify TFs with strong causal effects in specific species while also extending the identified driver regulators to a broader range of genes, particularly downstream genes of transcription factors and related enzymes. This capability likely stems from these genes’ direct roles in essential biological processes.

Following the evaluation of CauFinder on embryonic stem cell datasets, we expanded our analysis to include The Cancer Genome Atlas (TCGA) bulk data, which encompassing a broad array of cancer types (Fig. 2i). This transition allowed us to explore CauFinder’s effectiveness in a distinct biological context, specifically in identifying cancer drivers across diverse tumor types. We analyzed paired cancer and adjacent normal samples from eleven different cancer types within the TCGA dataset^42^, totaling over 1000 samples. For validation, we utilized a robust set of ground truth data sourced from the DisGeNET database, which provides a comprehensive collection of genes and variants associated with human diseases. To evaluate the precision and significance of CauFinder’s predictions, we measured the overlap between the drivers identified by CauFinder and the established pathogenic genes documented in DisGeNET. Our results indicate that CauFinder excels in identifying key cancer driver genes with high precision (Fig. 2j). In particular, the precision of CauFinder’s predictions increased with the number of ground truth genes available. This suggests that lower precision observed in certain cancers could be attributed to the limited number of known pathogenic genes, implying that CauFinder might uncover new potential cancer targets where ground truth data are sparse.

To further substantiate the performance of CauFinder, we conducted a comparative analysis with Mendelian Randomization^43^ (MR), which utilizes genetic variants as instrumental variables to infer causal relations in cancer biology. Our analysis focused on four cancer types, selected based on the availability of relevant SNP data. In each case, CauFinder’s precision outperformed MR, suggesting that CauFinder may offer more accurate identification of cancer drivers than genetic association-based methods alone (Fig. 2k). Further, CauFinder achieved these results without relying on SNP information, demonstrating its robust performance. Additionally, we explored the commonality of drivers across different cancer types identified by CauFinder. Notable among these are genes like PLK1 and RRM2, which are frequently overexpressed in various cancers, driving tumorigenesis and progression, and are therefore crucial targets for cancer therapy^44–50^. This analysis not only validates the effectiveness of CauFinder in detecting known drivers but also highlights its potential to uncover new drivers across cancers (Fig. 2l).

These findings from the TCGA dataset further confirm CauFinder’s robustness and versatility in handling complex genomic data from different biological contexts. By providing valuable insights into the mechanisms of cancer development and progression, this comprehensive evaluation highlights CauFinder’s potential to significantly impact genomic research. It facilitates the discovery of novel therapeutic targets and enhances our understanding of disease mechanisms.

### Identifying causal regulatory mechanisms and reversing the transdifferentiation from lung adenocarcinoma to squamous cell carcinoma

Human lung adenosquamous carcinoma (LUAS), accounting for approximately 0.7% to 11.4% of non-small cell lung cancer (NSCLC), is a distinct subtype characterized by greater plasticity, heightened malignancy, increased drug resistance, and poorer prognosis compared to lung adenocarcinoma (LUAD) and lung squamous cell carcinoma (LUSC)^51–58^. A recent study, utilizing the largest LUAS dataset to date, has demonstrated that the development of LUAS occurs through the transdifferentiation from lung adenocarcinoma to squamous cell carcinoma^59^. Researchers classified the LUAS samples into three categories based on cancer marker genes: the TRU-like subtype (LUAD-like), the inflammatory subtype, and the basal-like subtype (LUSC-like). Therefore, utilizing the dataset derived from the aforementioned study, we aim to investigate the capability of CauFinder in identifying causative driver factors/genes that are instrumental in the transdifferentiation from lung adenocarcinoma to squamous cell carcinoma.

We define samples in lung adenosquamous carcinoma with LUAD characteristics as the starting point of the state transition (state 0) and those with LUSC characteristics as the endpoint (state 1). Based on this definition, we utilize CauFinder to identify a group of driver factors with causal effects on adeno-to-squamous transdifferentiation (AST) (Fig. 3a, Supplementary Fig. 9a-b). We observed that TP63, SOX2, NKX2-1, and FOXA2, which were confirmed in previous studies, are present among these causal driver factors^59^. Additionally, this group of causal driver factors is significantly enriched in pathways associated with squamous cell carcinoma and adenocarcinoma, such as keratinocyte differentiation, epidermis development, epidermal cell differentiation, skin development, gland development, and keratinization (Fig. 3b). To highlight the characteristics of the LUAS samples, we performed UMAP dimensionality reduction and clustering (Fig. 3c, d). The dimensionality reduction results reveal that LUAS samples exhibit a trajectory distribution, with LUAD as the initial state and LUSC branching into two trajectories, both leading to LUSC as the endpoint, consistent with previous findings^59^. We inspected controllability scores for both the natural process (transition from LUAD to LUSC) and the reversal process (transition from LUSC to LUAD) on simulated state transitions manifold for all LUAS samples (Fig. 3c). All LUAD samples (transition from LUAD to LUSC) showed high and similar scores, whereas LUSC samples (transition from LUSC to LUAD) exhibited high scores only in one trajectory, indicating different transition capabilities. One LUSC branch had consistently high scores, possibly indicating a higher propensity to revert to LUAD. To further clarify the sample characteristics, we performed clustering on the samples (Fig. 3d), where LUAD and LUSC samples were grouped into two categories respectively (Supplementary Fig. 9d). Notably, one LUSC sample was clustered with the LUAD samples, likely due to its RNA expression pattern being very similar to LUAD, and this sample also had the highest state transition score (Fig. 3d). We further compared the difference in state transition scores between clusters, finding that each of LUAD and LUSC had a cluster with significantly higher state transition scores than other clusters (Supplementary Fig. 9e). This indicates the presence of an intermediate state prone to cause state transitions during the LUAS transformation process, which can be captured by CauFinder, consistent with the previous research findings^59^. Further, we observed the expression of several driver genes identified and validated by CauFinder (Fig. 3e), noting that they exhibited expression patterns similar to state transition scores, with each showing distinct variations. Subsequently, we extended the CauFinder model to include a functionality that simulates state transition trends by manipulating key driver factors. Utilizing this functionality, we employed samples in intermediate states and manipulated two key driver factors, TP63 and SOX2, which were previously validated through experimental studies. We specifically focused on three LUSC samples and observed that these samples indeed reversed from a squamous cell carcinoma-like state back to an adenocarcinoma state (Fig. 3f). We found that samples with higher scores (67 and 45) more easily transitioned from LUSC to LUAD compared to sample 53.

**Fig 3:**
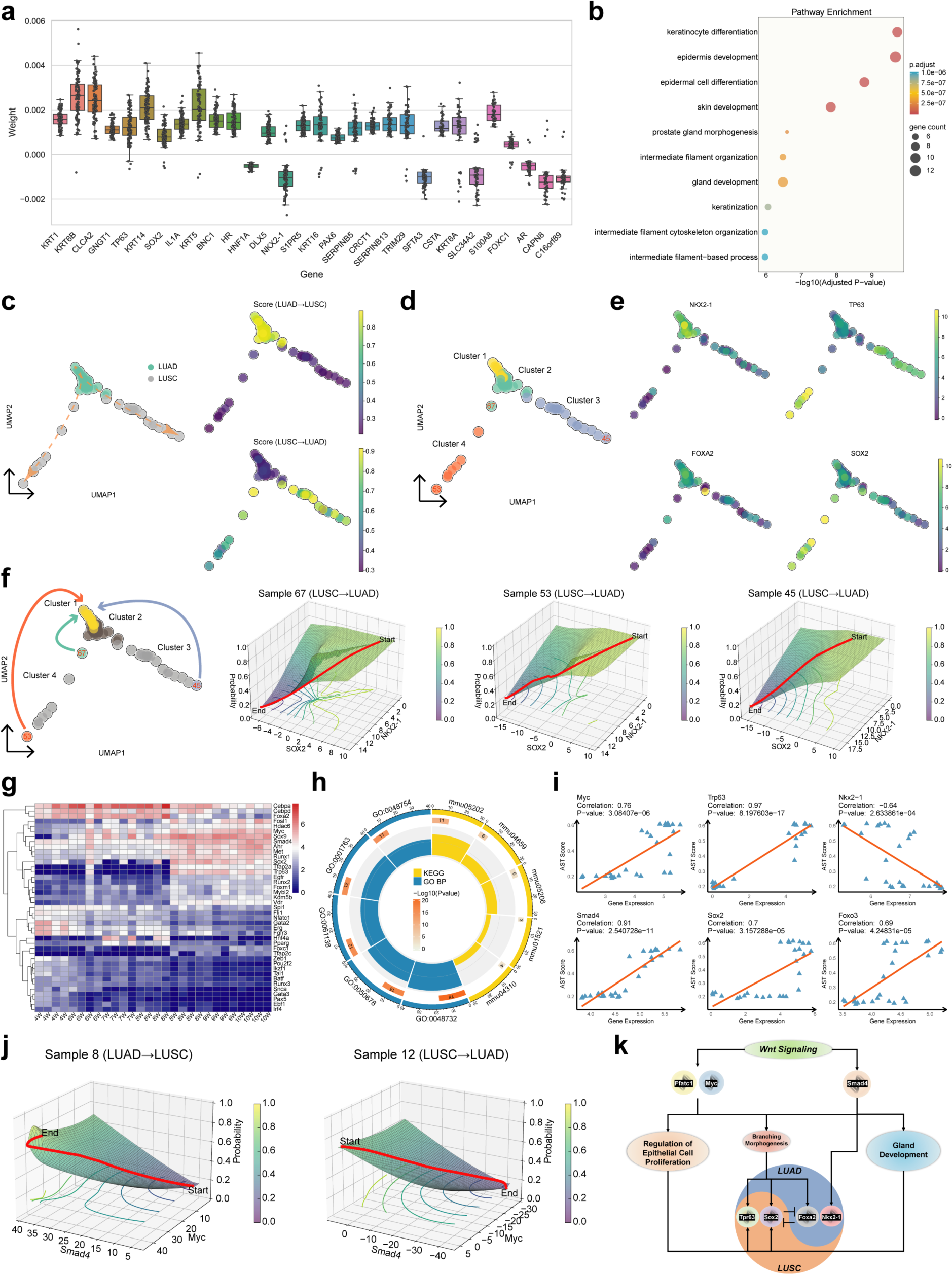
Identifying causal drivers igniting the transdifferentiation from lung adenocarcinoma to squamous cell carcinoma. **a**, Boxplots with individual data points showing the distribution of causal weight scores for the top 30 genes. These genes were selected based on their frequency of occurrence across 100 runs. The horizontal axis lists the genes, and the vertical axis represents the causal weight scores derived from the CauFinder model. **b**, Pathway enrichment bubble plot of the top 30 genes. The vertical axis lists the enriched pathways, while the horizontal axis shows the -log10(?) transformed adjusted p-values. Bubble colors indicate the adjusted p-values, with blue representing lower significance and red representing higher significance. Bubble size reflects the number of genes in each pathway, with larger bubbles representing more genes. **c**, Uniform manifold approximation and projection (UMAP) plot of RNA-seq data revealing the trajectory of phenotypic transitions from lung adenocarcinoma (LUAD) to lung squamous cell carcinoma (LUSC). Each node represents one sample, with colors indicating the tumor condition of samples (left) or representing the state transition score calculated by CauFinder: from LUAD to LUSC (top right) and from LUSC to LUAD (bottom right). **d**, UMAP plot showing the clustering results of samples using the Leiden algorithm with a resolution of 0.25. **e**, The expression patterns of four experimentally validated key genes identified by CauFinder are shown across the samples. These four genes exhibit distinct expression profiles in LUAD and LUSC, as well as along two trajectories within LUSC. The samples are distributed on the same UMAP coordinates as in (**d**), with each point representing a single sample. **f**, Computational simulation of the state transition for selected samples via control of causal genes. The leftmost panel shows the selection of three samples with a predominant LUSC state, which are controlled to transition towards LUAD. The middle and right panels depict the transitions of these three samples along distinct trajectories: Sample 67 (middle-left), Sample 53 (middle-right), and Sample 45 (rightmost). These panels focus on experimentally validated causal genes identified by CauFinder. Each panel illustrates the transformation of state probability during gene pair expression changes, where 0 indicates proximity to LUAD and 1 indicates proximity to LUSC. **g**, Heatmap of driver gene expression during the lung adeno-to-squamous transdifferentiation (AST) in mouse models. The horizontalaxis represents the weeks post Ad-Cre nasal inhalation, indicating the time points of 4W, 6W, 7W, 8W, 9W, and 10W. The verticalaxis displays driver genes with causal relations to AST, identified by CauFinder based on transcriptome data from mouse models. **h**, Circos plot of KEGG and GO pathway enrichment analysis for genes causally related to AST. The plot visualizes the top 10 most significant pathways. Each pathway is represented as a sector, with sector size proportional to the number of genes inputted into the database. Yellow sectors indicate KEGG pathways, while blue sectors indicate GO pathways. From the outer to the inner rings: the first ring outside the plot shows the pathway ID, the second ring shows the number of genes associated with each pathway, with colors representing -log10 P-values, and the innermost ring shows the GeneRatio (the ratio of enriched genes to the total number of input genes). **i**, Visualization of Pearson correlation coefficients for gene expression and the state scores of each AST sample. The horizontal axis shows gene expression levels, and the vertical axis displays samples state scores provided by CauFinder. **j**, Computational simulation of the state transitions (between LUAD and LUSC) via control of Myc and Smad4. The left panel shows the transition from LUAD to LUSC in a 7W sample, and the right panel shows the transition from LUSC to LUAD in a 9W sample. **k**, Conceptual model of AST mechanism based on CauFinder results. This model illustrates the role of Wnt signaling pathway inhibition in the AST process. Key genes such as Myc, Smad4, and Nfatc1 are shown impacting three critical pathways: regulation of epithelial cell proliferation, branching morphogenesis, and gland development. Dysregulation in these pathways contributes to the onset of AST.

This finding validates the capability of CauFinder in identifying driver factors with causal effects on AST, and demonstrates its potential in reversing squamous cell carcinoma-like states back to adenocarcinoma states. This suggests that these genes may be key factors influencing the LUAS transformation process. Through above analysis with CauFinder, we successfully demonstrated the reversal of squamous cell carcinoma back to adenocarcinoma. The identification of these driver factors not only highlights the potential to prevent the transition from adenocarcinoma to squamous cell carcinoma but also underscores the significant clinical implications of reversing squamous cell carcinoma to adenocarcinoma.

Having demonstrated the capability of CauFinder in identifying and reversing the transdifferentiation process, we next aimed to further expand upon the previously identified mechanisms contributing to this transition. Previous studies have established that the steady state between LUAD and LUSC is regulated by the Wnt signaling pathway, with its inhibition leading to lung adeno-to-AST^60^. However, the specific causal regulatory mechanisms remain poorly understood. We utilized CauFinder to analyze data from an established mouse model of AST previously characterized by other researchers^60^. In this model, mice were treated with Ad-Cre and classified based on the post-administration weeks: 4, 6, and 7 weeks (4W, 6W, and 7W) were defined as lung adenocarcinoma (LUAD) stages. The onset of AST was observed beginning in the 8th week (8W), which we designated as the intermediate AST state, progressing to lung squamous cell carcinoma (LUSC) by weeks 9 and 10 (9W and 10W)^60^.

Under the CauFinder framework, we defined and computed the state transition process with early adenocarcinoma stages (4W, 6W, and 7W) as the starting point (state 0) and late-stage squamous cell carcinoma (9W and 10W) as the endpoint (state 1). Additionally, we defined and computed the reverse process, with late-stage SCC as the starting points and adenocarcinoma stages as the endpoints. The intermediate state (8W) served as the validation dataset for CauFinder. Utilizing this framework, we identified a set of genes causally implicated in adeno-to-squamous transdifferentiation (AST) (Fig. 3g). Notably, this gene set included Foxa2, Sox2, and Trp63, which have been validated in the prior studies as being critical to this transdifferentiation process^59,60^. Pathway enrichment analysis was conducted on these identified genes, utilizing the GO database. The results highlighted that the most significant pathways were those related to gland development (GO:0048732), regulation of epithelial proliferation (GO:0050678), and branching morphogenesis (GO:0061138, GO:0001763, and GO:0048754) (Fig. 3h). These pathways are critically associated with the formation and progression of adenocarcinoma and squamous cell carcinoma. Furthermore, our analysis based on KEGG pathways revealed that this group of genes was significantly enriched in the Wnt signaling pathway (mmu04310), corroborating findings from previous the studies (Fig. 3h).

To elucidate potential new mechanisms, we next examined the distribution of causative genes within each significantly enriched pathway. Notably, previously validated genes such as Trp63, Sox2, and Foxa2 were present in pathways related to gland development, regulation of epithelial proliferation, and branching morphogenesis. Interestingly, genes within the Wnt signaling pathway predominantly co-localized with these pathways: Myc was involved in both the regulation of epithelial proliferation and branching morphogenesis; Smad4 was implicated in both branching morphogenesis and gland development; Nfatc1 was also linked to regulation of epithelial proliferation and branching morphogenesis. Although Nkx2-1, discussed earlier, was not identified by CauFinder as a causal gene, the identification of Smad4 by CauFinder as an upstream regulator with inhibitory effects^61^ suggests that Nkx2-1 may have a significant role.

Furthermore, to uncover the associations between samples, we conducted PCA analysis on all LUAD and LUSC samples collected at different time points. In the visualization of samples on PC1 and PC2, all samples are ordered by time (Supplementary Fig. 9f). The tipping point identified by the DNB method in the previous research^60^ also exhibits consistent behavior in state transition scores. Samples located at the tipping point have higher state transition scores, indicating a greater likelihood of state changes (Supplementary Fig. 9g, h). This represents a critical transition phase in the progression from LUAD to LUSC, consistent with the previous conclusions and demonstrating the usability and accuracy of CauFinder in simulating state transitions. Subsequently, to elucidate the state transition process in detail, we selected the most prominent samples based on state transition scores from LUSC and LUAD for further analysis (Supplementary Fig. 9i). Focusing on differentially upregulated genes Myc and Smad4, our analysis of Pearson correlation coefficients between gene expression and AST state scores defined by CauFinder (Fig. 3i), along with our computational simulations controlling key causal genes, indicated that managing Smad4 and Myc influences AST transitions comparably to previously validated genes (Fig. 3j, Supplementary Fig. 10). Additionally, we found that controlling Smad4 and Myc in these samples also demonstrated the potential to reverse AST (Fig. 3j). Therefore, Smad4 and Myc may represent genes of heightened interest to biological researchers. Additionally, since the pathway enrichment was based on causal genes, gland development, regulation of epithelial proliferation, and branching morphogenesis were more significant than the Wnt signaling pathway, suggesting that these three pathways may have a more direct causal relation with AST. Thus, the Wnt pathway is likely maintaining the AST steady state by modulating these three critical pathways through key causal genes (Fig. 4k). In this study, we employed the CauFinder algorithm not only to validate these earlier findings but also to reverse the transdifferentiation from adenocarcinoma to squamous cell carcinoma in mice based on newly identified causal genes. Additionally, we identified new regulatory mechanisms involved in AST.

**Fig 4:**
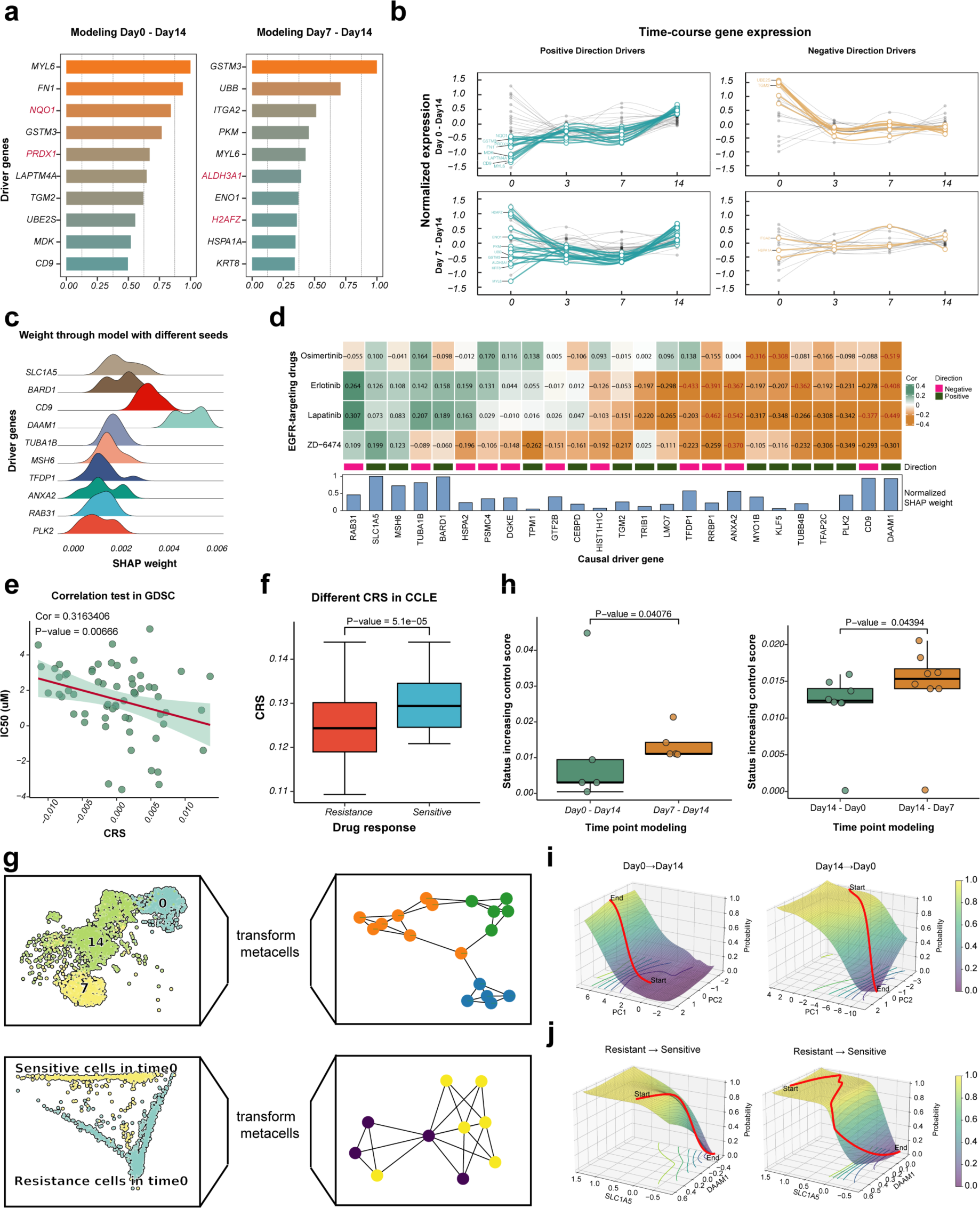
Characterizing causal gene expression programs in cycling persister cells arising and in response to different EGFR-targeting drugs. **a,** Top ten causal drivers identified by CauFinder, based on lineage barcode tracing of progenitor cells of cycling persister cells sequenced on day 0 (left) and day 7 (right), paired with corresponding cycling persister cells on day 14 as the input. The drivers are ranked by their normalized SHAP weights. Genes involved in the ROS and FAM pathways are highlighted in red. **b,** Time-course expression of causal driver genes identified by CauFinder for two different paired inputs. Expression is represented as the normalized average expression of the corresponding genes across all lung cancer cells in each time category. Causal drivers are grouped by the direction of their weights from the CauFinder output, with the top ten drivers highlighted. Drivers with positive weights are shown in blue, while those with negative weights are shown in orange (Results from day 0 to day 14 input, upper; results from day 7 to day 14 input, lower). **c,** Top ten causal drivers identified by CauFinder, based on results from using sensitive cells paired with resistant cells as input. The density estimation of SHAP weights output by CauFinder is shown for each causal driver, with different seeds used for robustness. The drivers are ranked by their SHAP weights. **d,** Heatmap showing the Pearson correlation between drug response IC50 values of patients treated with four EGFR-targeting drugs in GDSC and CCLE databases and the expression of each causal driver gene across patient samples. Correlation pairs with a p-value less than 0.05 are highlighted. The color bar indicates the direction of the causal drivers’ weights as determined by CauFinder. The bar plot at the bottom represents the normalized SHAP weights of the corresponding causal drivers. **e,** Correlation between causal drug response score (CRS) and IC50 values (horizontal and vertical axes) in patients (dots) from the GDSC database. **f,** Boxplots showing the distribution of CRS of patients treated with three EGFR-targeting drugs in the CCLE database across two drug response groups. P-values were obtained from the T-test. The middle horizontal line represents the median value. Each box spans the lower quartile to the upper quartile. The whiskers indicate the minimum and maximum values within 1.5 times the interquartile range (IQR). **g,** UMAP plots of original cells from time-course pairs (top) and drug response pairs (bottom) with a pseudo-trajectory network of transformed pseudo-cells used as input for the CauFinder state transition function. Both original and pseudo-cells are grouped by time. Edges in the pseudo-cell network are inferred by the Partition-based graph abstraction (PAGA). **h,** Pseudo-cell state transition scores generated from two different start state cell inputs, with the same end state cell input. The p-values between the two groups are shown on the top (the Kolmogorov-Smirnov test). Forward state transitions (using the end state as target) are shown on the left, and reverse state transitions (using the start state as target) are shown on the right. **i,** Computational simulation of state transitions for different samples from the time-course pairs. Principal components (PCs) identified through principal component analysis (PCA) of all drivers. The X-axis and Y-axis represent the principal components (PCs) of these controlling drivers, while the Z-axis indicates the state probability, where values closer to 0 represent the defined starting point and values closer to 1 represent the defined endpoint. **j,** Computational simulation of state transitions for different samples from the drug response pairs regulated by DAMM1 and SLC1A5. The X-axis and Y-axis represent the expression levels of DAMM1 and SLC1A5, respectively, while Z-axis indicates the state probability, analogous to (**i**).

### CauFinder identifies regulatory mechanisms of lung cancer cycling cancer persister cells and guides improvement of drug response

To further validate the applicability of CauFinder on real data and demonstrate its robustness and versatility across temporal dimensions, we collected a dataset with multiple time points from a previous study on cancer. The study focused on PC9 lung cancer cells, particularly a class of drug-resistant, continuously proliferating cells known as cycling cancer persister cells. Using the Watermelon system^62^ for lineage tracing, the dataset allows tracking of lineage relations among cells sequenced at different time points at the single-cell level.

First, we generated three sets of paired states based on known information: cycling cancer persister cells sequenced on day 14 paired with their ancestral persister cells from the same lineage on day 0 (day0-day14), cycling cancer persister cells sequenced on day 14 paired with their ancestral persister cells from the same lineage on day 7 (day7-day14), and cycling cancer persister cells sequenced on day 14 paired with non-cycling cancer persister cells sequenced simultaneously (cycling-noncycling). These three sets of pairs were used as inputs for CauFinder. CauFinder identifies causal drivers between the input states and assigns weights to each driver. Among the top-weighted causal drivers identified from the three different inputs, we consistently found genes related to reactive oxygen species (ROS) and fatty acid metabolism (FAM) from previous studies (Fig. 4a, Supplementary Fig. 11a), we compared all similar sets of four input pairs and observed that the drivers identified by CauFinder exhibit a certain level of consistency within the same biological processes (Supplementary Fig. 12b). This consistency, even with different random seeds, demonstrates the robustness of CauFinder (Supplementary Fig. 12a). In particular, ROS-related genes (NQO1 and PRDX1) and FAM-related genes (ALDH3A1 and H2AFZ) had relatively higher weights in the day0-day14 paired model and the day7-day14 paired model (Fig. 4a). We hypothesize that this result stems from the redox balance being regulated early in the drug-stress process, while FAO plays a role in establishing the proliferative persister state later. To further test this hypothesis, we examined the weight distribution of causal drivers within the Hallmark gene set. In both the day0-day14 paired model and the day7-day14 paired model, ROS and FAM-related causal drivers showed higher overall weights compared to other gene sets (Supplementary Fig. 11b). Gene set enrichment analysis (GSEA) using the Kyoto Encyclopedia of Genes and Genomes (KEGG) highlighted pathways potentially influencing the regulation of proliferation in cycling cancer persister cells (Supplementary Fig. 11c-f).

Next, we analyzed the temporal expression patterns of causal drivers identified from the two paired inputs (day0-day14 and day7-day14). We used the average expression of the corresponding genes across all cancer cells sequenced at each time point to represent gene expression levels (Fig. 4b). The causal drivers from both inputs exhibited consistent temporal expression patterns, corresponding to the weight direction in the CauFinder results. Causal drivers with positive weights showed increased expression towards the target state (day14), while those with negative weights showed the opposite trend. The two different initial states (day0 or day7) resulted in distinct expression patterns for causal drivers, with the day0 input showing a consistent expression pattern across all four time points, while the day7 input showed consistency primarily between day7 and day14. These findings suggest that CauFinder can accurately uncover specific features and robustly identify causal drivers, representing unique connections between initial and target states.

CauFinder’s capability to explore sample state transitions can also be applied to single-cell data. To address the sparsity and high number of cells in single-cell data, we introduced pseudo-cells to represent the original single-cell data. We grouped the original single-cell data by initial and target state labels, re-clustered within groups, and used the average expression of cell subpopulations to represent the groups. This conversion reduced the computational load and avoided filtering effective results from a large output. We transformed the three-state data (day0-day7-day14) into pseudo-cells and combined them for dimensionality reduction. We obtained five pseudo-cells representing the initial states (day0 and day7) and eight pseudo-cells representing the common target state (day14) (Fig. 4g, Supplementary Fig. 12c).

We tested CauFinder output results in two transition directions: the forward state transition, with day14 as the endpoint, and the reverse state transition, with day14 as the starting point. Using the defined status control score to evaluate state transitions, we compared the status control scores of different pseudo-cells in both transition directions (Fig. 4h). Regardless of the transition direction, models with day7 as the starting point had significantly higher state transition scores than those with day0 as the starting point (P-value = 0.0407 for increasing, P-value = 0.0439 for decreasing, by the Kolmogorov-Smirnov test). Computational simulations of state transitions for each pseudo-cell showed different trends. All pseudo-cells derived from day 0 and day 7 cells successfully underwent the expected forward state transition to day 14, consistent with biological processes. In addition, some pseudo-cells with high state transition scores, derived from day 14 cycling cells exhibited a strong reverse state transition to earlier stages (Fig. 4h). This suggests that CauFinder has potential applications in guiding the regulation of reverse state transitions in biological processes (Fig. 4i). Pseudo-cells in day0 exhibited smaller probability changes during forward state transitions compared to those in day7. Notably, pseudo-cell 4 in day0 had a higher status control score compared to other pseudo-cells in the same group (the state control score = 0.044794) (Fig. 4h). We further examined the differential expression between pseudo-cell 4 and other pseudo-cells (Supplementary Fig. 12d-e), identifying genes such as POLR3A and COX19, which are related to cancer progression or drug resistance and may be involved in the regulation of cycling persister cells. These findings highlight the critical roles of ROS and FAM in the regulation of cycling persister cells, demonstrating capability of CauFinder to capture key factors in complex biological processes and guiding the regulation of reverse state transitions in biological processes.

We further partitioned the same dataset according to another biologically significant feature with the broader medical value and application potential: tumor cell drug resistance. We categorized the dataset into two groups based on cell sensitivity and resistance to Osimertinib. Based on the causal drivers identified by CauFinder using different seeds, we ranked the top ten causal drivers by weight, which included several genes associated with cancer and tumor resistance, such as CD9, BARD1, and SLC1A5 (Fig. 4c). We initially explored the correlation between the expression of all causal drivers and the IC50 of four EGFR-targeting drugs in the CCLE and GDSC datasets (Fig. 4d). Significant correlations were found for TFDP1, RRBP1, ANXA2, DAAM1, CD9, MYO1B, KLF5, and TUBB4B with the sensitivity of one or more drugs (P-value < 0.05, by the Pearson correlation). This indicates that CauFinder can capture a variety of genes related to drug sensitivity. However, some causal drivers did not show significant correlations between single gene expression and patient IC50 values. We believe that causal drivers are not independent but function collectively in biological processes.

Therefore, we weighted gene expression by the weights of the causal drivers from CauFinder and summed them according to their weight direction to obtain a causal drug response score (CRS) representing the sample drug response. In the GDSC dataset, CRS showed a significant correlation with IC50 (P-value = 0.006, by the Pearson correlation) (Fig. 4e). When samples were divided into two equal groups based on the median CRS (High and Low), the IC50 values between the two groups showed a significant difference (P-value = 0.021, by the T-test) (Supplementary Fig. 12f). Similarly, in the CCLE dataset, patients were divided into two groups based on drug sensitivity labels (IC50 > 8 marked as resistant). The CRS of the sensitive group was significantly higher than that of the drug-resistant group (Fig. 4f), and also CRS showed a significant correlation with IC50 (P-value = 0.006, by the Pearson correlation) (Supplementary Fig. 12g), indicating that CRS can effectively distinguish whether clinical patients are resistant to EGFR-targeting drugs. Independently, we evaluated whether the causal drug response score could be used as a yardstick for survival risk stratification. A significant improvement in overall survival was observed in patients with a low causal drug response score (Supplementary Fig. 12h). A multivariate Cox regression analysis yielded a likelihood ratio test P-value = 0.02.

To explore the potential for sample state transitions in single-cell datasets as input for the model, we used the same method as before to convert the original single-cell expression matrix into a reduced number of pseudo-cell expressions to reduce sample size (Fig. 4g, Supplementary Fig. 12i), highlighting the common characteristics expressed by similar samples in the single-cell data. We generated four pseudo-cells for the sensitive state and six for the resistant state. It was observed that sensitive state cell-0 and cell-3 were more closely linked to the pseudo-cells of the resistant state, while resistant state cell1 was more closely linked to the pseudo-cells of the sensitive state.

We examined specific simulated state transition processes (Fig. 4j, Supplementary Fig. 13a-f), where most pseudo-cells achieved overall state transitions. Using some pseudo-cells as examples, causal drivers in drug response identified by CauFinder (DAAM1 and SLC1A5) were used as operative genes, efficiently achieving simulated transitions and altering cell drug sensitivity. Although CauFinder can simulate state transitions for both drug-resistant and drug-sensitive pseudo-cells, a comparison reveals that while the biologically significant forward process (tumor enhanced tumor drug resistance) can be straightforwardly modeled, reducing tumor cell drug resistance, though possible, results in more complex and difficult-to-regulate transition paths (Fig. 4j). This aligns with the real-world observation that tumor cells are more prone to spontaneously develop drug resistance rather than spontaneously reducing it.

CauFinder effectively identifies potential gene therapy strategies to enhance sample drug sensitivity by modulating the expression levels of genes identified as causal drivers. This approach can improve the efficacy of specific drug treatments, offering new methods for treating drug-resistant cancers and supporting cancer medication.

### CauFinder defines key intermediate cell states and interaction mechanisms in primary colorectal cancer (P1) and paired liver metastasis (LM1)

To validate the potential of CauFinder in spatial transcriptomics analysis, we selected a dataset of primary colorectal cancer (P1) and paired liver metastasis (LM1) from a recent study^63^ (Fig. 5a, c). For the primary colorectal cancer sample P1, we initially applied the MCGAE algorithm to cluster the spatial transcriptomics data. By combining the clustering results with manual annotation, we divided the data and retained only the tumor tissue and normal tissue sections as input for training the CauFinder model (Fig. 5a). Similarly, we divided and filtered the paired liver metastasis sample LM1. However, unlike the P1 sample, we selected only the two cell clusters at the tumor-liver interface (cluster 1 and cluster 6) to represent tumor tissue and liver tissue, respectively, as input (Fig. 5c). All samples were trained ten times using different random seeds.

**Fig 5:**
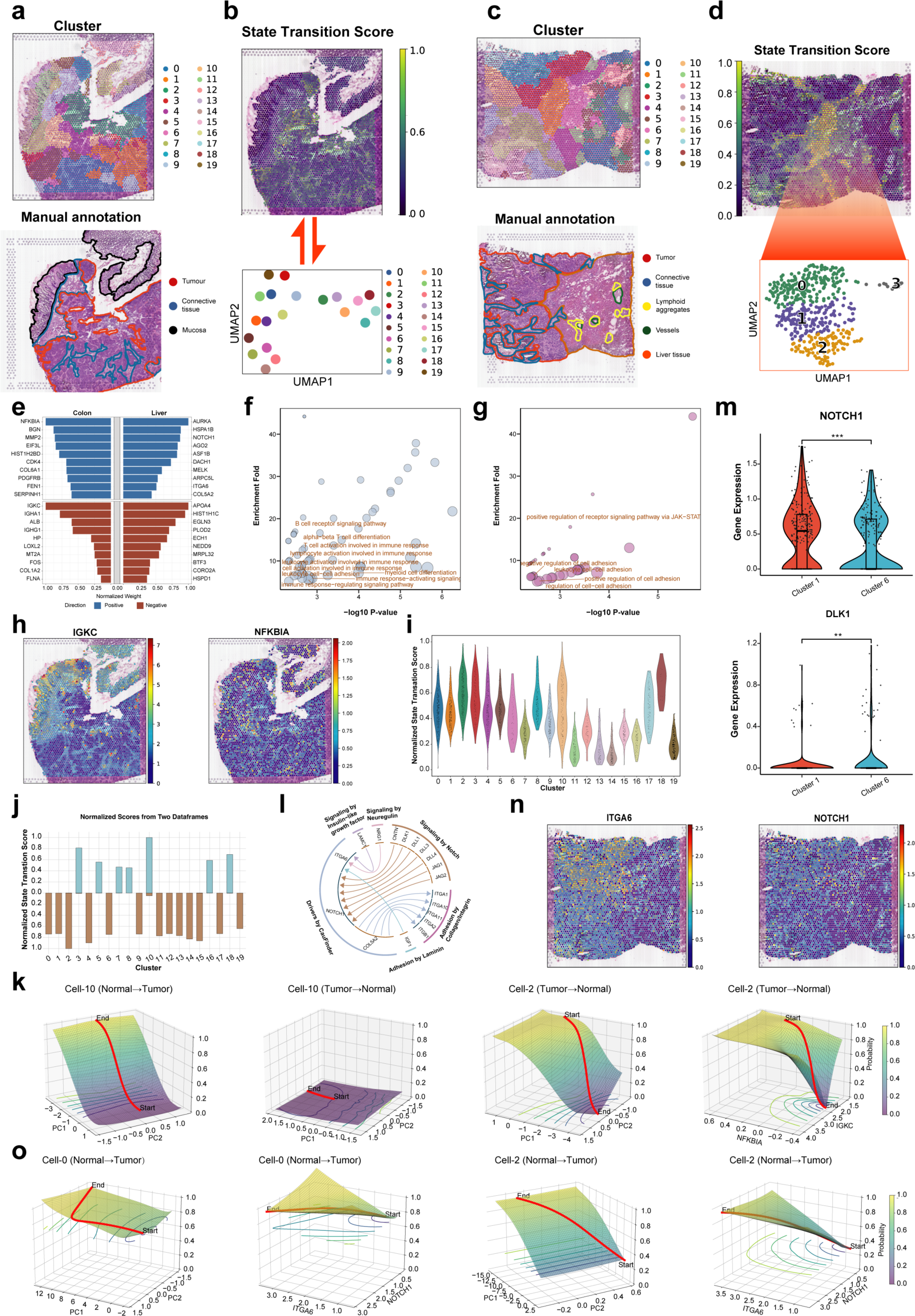
Identification of causal driver genes and intercellular interactions in colorectal cancer at the spatial transcriptomics level. **a,** Hematoxylin and eosin (H&E) plot of P1 tissue (top) with manual annotation of three regions: tumor (red), connective tissue (blue), and mucosa (black). Spatial clustering by MCGAE on P1 tissue (bottom). **b,** Spatial distribution plot of cell state transition scores of P1 tissue (top). Cells are re-clustered and converted to pseudo-cells, which are then used as input for the CauFinder state transition function (bottom). **c,** Hematoxylin and eosin (H&E) plot of LM1 tissue (top) with manual annotation of five regions: tumor (red), connective tissue (blue), lymphoid aggregates (yellow), vessels (green), and liver tissue (orange). Spatial clustering by MCGAE on LM1 tissue (bottom). **d,** Spatial distribution plot of cell state transition scores of LM1 tissue (top). Cells are re-clustered and converted to pseudo-cells, which are then used as input for the CauFinder state transition function (bottom). **e,** Top ten causal drivers with positive and negative weights defined by CauFinder in P1 tissue (left) and LM1 tissue (right). Bar length indicates the absolute value of the weights, with color representing the direction: positive (blue) and negative (red). **f,** Bubble plot of GO BP enrichment analysis using all identified causal driver genes in P1 tissue, with immune-related pathways labeled. The size of each point represents the number of genes enriched in the pathway, the horizontal axis shows the log-transformed P-value, and the vertical axis indicates the enrichment fold ((enrichment fold = GeneRatio/ BgRatio, provided by clusterProfiler). **g,** Bubble plot of GO BP enrichment analysis using all identified causal driver genes in LM1 tissue, presented in the same manner as in (**f**). **h,** Spatial expression of IGKC and NFKBIA genes in P1 tissue. **i,** Violin plot showing cell state transition scores among different clusters, determined by clustering results, with each cluster represented by a different color. **j,** Cell state transition scores of pseudo-cells, converted from single cells by cluster, simulating transition to two states (tumor: 1, normal tissue: 0). Blue indicates transition to the tumor state and red indicates transition to the normal tissue state. **k,** Computational simulation of state transitions in a subset of pseudo-cells from P1 tissue. Principal components (PCs) identified through principal component analysis (PCA) of all drivers. The X-axis and Y-axis represent the principal components (PCs) of these controlling drivers, while the Z-axis indicates the state probability, where values closer to 0 represent the defined starting point and values closer to 1 represent the defined endpoint. **l,** Circos plot of ligand-receptor interactions involving causal drivers in LM1 tissue, with each arrow pointing from ligand to receptor. The outermost text represents different ligand-receptor sets as described in CellPhoneDB. **m,** Expression of NOTCH1 gene and its ligand DLK1 at the boundary between two clusters of tumor and normal tissue. **n,** Spatial expression of NOTCH1 and ITGA6 in LM1 tissue. **o,** Computational simulation of state transitions in a subset of pseudo-cells at the tumor-liver tissue interface in LM1 tissue. The X-axis and Y-axis represent the same variables as in (**k**).

We subsequently identified a set of causal drivers determined by CauFinder, retaining those found in at least five models with different random seeds to ensure accuracy. The top ten causal drivers were ranked by the absolute values of their weights, categorized into positive and negative weights (Fig. 5e). In the P1 sample, the results included immune-related genes (IGKC, NFKBIA, IGHG1), whereas in the LM1 sample, the results included cell ligand-receptor genes and surface adhesion-related genes (NOTCH1, COL5A2, HSPD1). Next, we performed biological pathway enrichment analyses of the causal drivers in P1 and LM1 (Gene Ontology: biological process, P < 0.01). The biological pathways enriched in the P1 sample included many immune-related pathways (Fig. 5f). Similarly, the biological pathways enriched in the LM1 sample included numerous cell adhesion-related pathways and signaling pathways (Fig. 5l). Moreover, mapping the causal drivers from the LM1 sample to the CellPhoneDB cell communication database revealed multiple interaction pairs. Among these, interaction pairs such as DLK1-NOTCH1 exhibited opposite differential expression in adjacent tumor and normal tissues, indicating potential reciprocal regulation (Fig. 5m, n).

These findings suggested that CauFinder’s results can preliminarily characterize the impact of immune responses in the transition of normal tissue to adjacent cancer tissue in primary colorectal cancer. In contrast, in liver metastasis, the cell-cell communication between colorectal cancer tissue and adjacent liver tissue may be a primary driver of liver metastasis progression. We used the causal drivers identified in the P1 and LM1 samples to independently simulate state transitions for all single cells in each sample. For any given cell, we considered both forward and reverse state transitions, selecting the direction with the higher state transition score as the determined transition direction for that cell. The resulting state transition scores were then spatially displayed (Fig. 5b, d).

In the P1 sample, we observed that cells with higher state transition scores were concentrated in the connective tissue region between the tumor and mucosa, as well as in a group of tumor cells adjacent to and connected with this connective tissue region. We examined the expression of two immune-related genes (IGKC and NFKBIA) among the causal drivers in the P1 sample (Fig. 5h). IGKC was highly expressed in the mucosa and connective tissue, while NFKBIA was highly expressed in the tumor tissue. We also noted that higher expression levels in the tumor cells adjacent to and connected with the connective tissue region were consistent with the state transition scores. We categorized all cells based on clustering results (Fig. 5i), finding that tumor cell clusters adjacent to the connective tissue region (clusters 2, 3, 8, 10) exhibited higher state transition scores, and some clusters corresponding to the connective tissue also showed higher state transition scores.

To mitigate the impact of excessive single-cell data, we converted the cells in the P1 sample into pseudo-cells, using the same processing method as for the cycling cancer persister cells (Fig. 5b). We performed simulated state transitions on pseudo-cells and recorded the state transition scores for each pseudo-cell (Fig. 5j). Among them, pseudo-cells like 5, 7, 8, 10, 16, and 18 exhibited a strong potential for transitioning to cancerous tissue, while other pseudo-cells showed a strong potential for transitioning to normal tissue. We used cell-2 and cell-10 as examples. Cell-10 could be regulated to a tumor state but not to a normal state (Fig. 5k). This may be because, in the P1 sample, the cells in Cluster 10, although located at the tumor-normal tissue interface and containing both normal and tumor cells, had fewer tumor cells and possibly represented early-stage tumor cells, retaining more characteristics of normal cells. Therefore, similar to pseudo-cells derived from normal cells like cell-18, they lacked the ability to transition back to their original state (Supplementary Fig. 14c). Cluster 2, as a type of boundary tumor cell, might also represent early-stage tumor cells but possessed more tumor characteristics, enabling pseudo-cells like cell-2 to undergo reverse state transitions to normal tissue, akin to other tumor cells (Fig. 5k, Supplementary Fig. 14a). This process suggests potential cancer treatment strategies. We also selected two gene combinations that have been confirmed in studies to have the potential for developing anti-tumor drugs^64^. Under these two gene combinations, we could achieve state transitions for many pseudo-cells, including cell-2, derived from tumor cells, transitioning them to normal tissue (Supplementary Fig. 14b). This demonstrates that CauFinder can effectively define the key factors in the spatial transition process between tumor and normal tissues and guide the reverse regulation of this process, providing potential targeted anti-tumor therapeutic strategies.

In the LM1 sample, we observed a similar pattern to that in the P1 sample, where cells at the interface between tumor tissue and normal tissue exhibited higher state transition scores. The region with the highest state transition scores largely overlapped with the cluster 1 identified through clustering (Fig. 5d). Notably, these cells were not labeled during CauFinder model training, indicating that this result was not influenced by the training process. We converted cells from this region and adjacent areas into pseudo-cells and further examined the specific state transition processes. We found that pseudo-cells derived from the adjacent region also exhibited differences in state transition processes (Fig. 5o). When we selected a set of potential genes (DLK1 and ITGA6) from the causal drivers, the pseudo-cells derived from cells located between tumor and liver tissue, despite some cells, such as cell 0, being identified by the model as having a state closer to that of tumor cells compared to other pseudo-cells, could all be regulated to transition to a tumor state. This suggests that CauFinder can also identify key driver genes in the tumor metastasis process and simulate their patterns of change, providing valuable insights for exploring specific biological mechanisms.

## Discussion

In this work, we introduce CauFinder, a sophisticated framework designed to accurately identify causal regulators of cell-state/phenotype transitions and further precisely steer such transitions, by integrating causal disentanglement modelling with nonlinear network control based only on the observed data. Understanding and manipulating cell states and phenotype transitions are crucial for advancing biological research and therapeutic strategies. Our framework is particularly valuable for disentangling complex causal relations within high-dimensional biological data.

CauFinder consists of two main components: a causal decoupling model and a network control mechanism. The causal decoupling model employs variational autoencoders to identify latent causal factors, separating them from spurious factors. We then utilize SHAP values and gradient information to trace these latent factors back to the original feature space, thereby identifying key causal features. The network control mechanism uses the Minimum

Feedback Vertex Set (MFVS) method, integrated with causal weights, to identify master regulators and guide effective interventions.

Our experiments demonstrated that CauFinder could effectively identify and manipulate causal features/genes to achieve desired phenotype transitions. By applying our framework to both simulated and real-world data, we observed significant improvements in intervention strategies compared to various traditional methods. The use of prior gene interaction networks enhanced the robustness of our model, and the staged training strategy ensured optimal performance across different loss terms. The controllability score provided a quantitative measure of the ease of controlling each sample, aiding in the evaluation of intervention effectiveness. In particular, CauFinder is able to not only reveal natural biological transition processes such as cell differentiation processes, LUAD to LUSC transdifferentiation (lung cancer), and drug-sensitive to resistance process, but also identify the causal regulators of many “reversal” biological transition processes such as LUSC to LUAD transdifferentiation, and drug resistance to drug sensitive processes.

Despite its strength, CauFinder has some limitations. One limitation is the potential impact of unobserved variables, which can confound causal inference and affect the accuracy of the identified causal features. However, as discussed in the Supplementary Note 6, if the observed data encapsulates all relevant causal information concerning y, the presence of some unobserved variables does not impact the results. This condition is generally met in high-dimensional feature spaces, such as scRNA-seq data. Further there are several avenues for future research that can enhance the capabilities of CauFinder. Extending the framework to handle multi-class phenotypes would broaden its applicability and allow for a more nuanced understanding of complex biological processes. Developing methods to directly integrate continuous variables without discretization would improve the theoretical robustness of our approach. Expanding the framework to address many-to-many relations in high-dimensional data could enhance its ability to model hierarchical controls within biological systems. Additionally, incorporating more precise network control algorithms, such as conflict multi-objective optimization techniques^65^, could further refine the intervention strategies and improve the overall effectiveness of our framework.

In conclusion, CauFinder represents a significant step forward in achieving cell state or phenotype transitions by integrating causal modelling and network control, based on the observed data. By addressing its limitations and exploring future directions, we can further enhance its utility and applicability in biological research and therapeutic interventions.

## Methods

### Causal decoupling/disentangling model

We begin with a dataset composed of a collection of samples *X* = [*x*_1_, … , *x*_n_]^*T*^, within ℝ^*n*×*p*^, where each sample *x*_*i*_ contains *p* features. Each sample *x_i_* is associated with a binary variable *y*_*i*_ ∈ {0, 1}, representing the phenotype or state. Our study aims to identify key features that significantly influence state or phenotype transitions through causal mechanisms. By isolating these features, we seek to understand and manipulate the underlying causal factors driving these transitions.

#### Structural causal model

In this subsection, we analyze the learning process from input features *x* to the response variable *y* from a causal perspective. Our approach begins by identifying potential causal factors in the latent space before pinpointing causal features in the original space. This strategy enables us to circumvent the complexities associated with the interdependencies in the input space. To this end, we construct a structural causal model (SCM)^18^ in Fig. 1b, where *x* = {*x*^*c*^, *x*^*s*^} represents features in the original space and *z* = { *z*^*c*^, *z*^*s*^} denotes features in the latent space, with *y* representing the corresponding state or phenotype. Here, *x*^*c*^ and *z*^*c*^ represent the causal components, while *x*^*s*^ and *z*^*s*^ signify the spurious or non-causal parts.

According to our causal hypothesis, when considering the causal relation between latent features and the output, a confounder arises from the data space. Specifically, when exploring the causal effects of *z*^*c*^ on *y*, we identify a backdoor pathway: *z*^*c*^ ← *x* → *z*^*s*^ → *y*, where *x* acts as a confounding variable between *z*^*c*^ and *y* . The presence of this confounder is a significant obstacle in causal quantification. Inspired by recent advancements in information theory^66,67^, we employ information flow (denoted as *I*(*z*^*c*^ → *y*) to quantify the causal impact of the latent factor *z*^*c*^on model prediction *y*. It enables us to accurately measure the causal information transfer from latent features to predicted outcomes. Ultimately, we implement this causal model using a dual autoencoder, with implementation details provided in Fig. 1c.

#### Dual variational autoencoder

Within the SCM framework, the dual variational autoencoder (DVAE) serves as the foundational mechanism, specifically designed to operationalize the model and critically extract and distinguish causal from spurious features within complex datasets. The model consists of two distinct branches: the causal branch and the spurious branch. In the causal branch, input features *x* are weighted via learnable weights *w*, resulting in *x^w^* = *x* ⊙ *w*, where ⊙ denotes element-wise multiplication. These weighted features are subsequently transformed into the causal latent space *z*^*c*^ by a causal encoder and then reconstructed by the decoder. In parallel, in the spurious branch, *x* is processed by complementary weights *w*= = 1 − *w*, forming *x^w^* = *x* ⊙ *w*=. It is then projected into the non-causal, or spurious latent space *z*^*s*^and reconstructed. The outputs from these two latent spaces are utilized to predict the sample’s state or phenotype *y* . The reconstruction process in both branches focuses on recovering features in the original space that are heavily weighted by their respective *w* and *w*= , ensuring that *z*^*c*^ and *z*^*s*^ collectively capture all the information from the original space, each with its respective emphasis.

Each VAE within the DVAE aims to maximize the Evidence Lower Bound (ELBO), which serves as a lower bound on the log likelihood of the real data distribution log *p*(*x*). The ELBO, central to the optimization of variational autoencoders, comprises two components: the reconstruction loss and the Kullback-Leibler (KL) divergence. The reconstruction loss ensures that the mapping from the latent space back to the data space is as accurate as possible, while the KL divergence measures the closeness of the encoded latent variable distribution to its prior distribution. Specifically, the ELBO loss is formulated as:

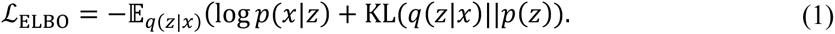

where KL(*q*(.)||*p*(.) is the Kullback-Leibler divergence between *q*(.) and *p*(.). Following the encoding process, the latent representations *z* are used to predict the state *y*. For this prediction task, the standard binary cross-entropy loss is employed:

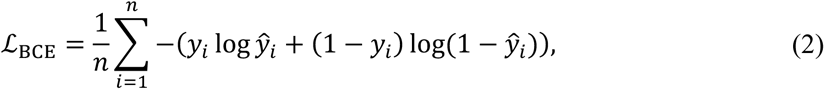

where *y_i_* is the true label of the *i-*th sample, and *y*L is the predicted probability of the *i*-th sample belonging to class 1.

The key to DVAE lies in leveraging the intrinsic structure of the data within the latent space to differentiate between causal and spurious factors. In subsequent sections, we will demonstrate how optimized quantification of causal information flow can achieve the decoupling of causal and spurious features in the latent space.

#### Causal information flow measurement

Identifying potential causal factors in the latent space offers two significant advantages: first, by harnessing the inherent properties of variational autoencoders (VAEs), the variables within the latent space z are mutually independent, allowing us to bypass the complex interdependencies in the input space; second, the backdoor paths associated with the latent variables in *z* are well-defined, providing a clear method to circumvent confounding effects and enabling an accurate measurement of causal information flow from latent features to predictions. The challenge lies in quantifying the causal effects of different aspects of the data within the latent space to identify the most causally significant components. As mentioned above, we use information flow *I*(*z*^*c*^ → *y*) to quantify the causal impact of *z*^*c*^ on *y* , which can be seen as the causal counterpart of mutual information *MI*(*z*^*c*^; *y*) . Our framework aims to maximize this information flow, thereby isolating the subset of latent variables *z*^*c*^ that represent causal features.

When deliberating on the causal relation between latent features (i.e., causal and spurious factors) and model predictions, *x* acts as a confounder. Ignoring *x* might lead to inaccurate estimations of causal features. To mitigate this, we employ the classic backdoor adjustment formula^23^, which is presented as follows:

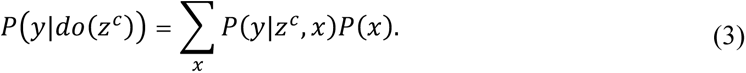

Equation (3) is crucial for bypassing confounding effects introduced by the input feature and for computing the information flow *I*(*z*^*c*^ → *y*), which is the causal counterpart of mutual information. Intuitively, this formula estimates the causal effect of *z*^*c*^ on *y* by considering different versions of *x* while keeping *z*^*c*^ fixed. In causal theory, this approach, known as do-calculus, effectively eliminates the influence of confounders by applying interventions to *z*^*c*^.

The information flow between the causal factors *z*^*c*^ and the prediction *y* can be computed as follows:

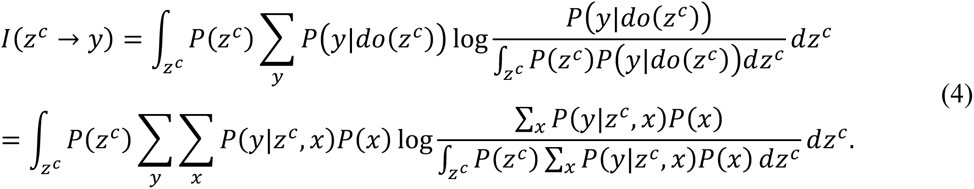

For a detailed derivation of *I*(*z*^*c*^ → *y*), see Supplementary Note 5. Our primary aim is to maximize the causal information flow measure from *z*^*c*^ to *y*. Therefore, the corresponding loss term for this objective is given by

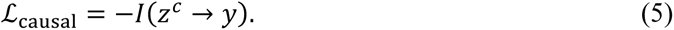

To ensure that *z*^*c*^captures the main information about *y*, we introduce a fidelity loss. The goal here is to ensure that the distribution of *y* calculated from *z*^*c*^is as close as possible to the distribution of *y* obtained from the entire latent space *z* . This ensures that our causal components indeed represent the main influence on the outcome. The fidelity loss can thus be expressed as follows:

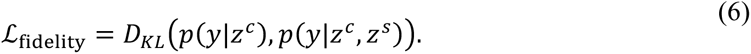

When considering the causal information flow measure *I*(*z*^*c*^ → *y*), we also examine the impact of unobserved variables on its definition. For detailed analysis and proofs, refer to Supplementary Note 6.

#### Identifying causal features in the original space

After decomposing the latent space features, we obtain the causal pathway *x* → *x*^*c*^ → *y* . While latent space features provide valuable insights, our primary interest lies in identifying causal features within the original space, denoted as *x*^*c*^. To achieve this, we employ SHapley Additive exPlanations (SHAP) values to determine the causal weights of individual features along the causal pathway. We compute the SHAP values to quantify each feature’s contribution to *z*^*c*^ and subsequently to *y*, thereby identifying the features that play pivotal roles in the causal pathway. Then, we utilize gradient calculations to ascertain the direction of influence of the original space features on state transitions. A positive gradient indicates that higher expression of the feature promotes the transition of *y* from state 0 to state 1, whereas a negative gradient indicates that higher expression inhibits this transition. This dual approach of SHAP values and gradient calculations allows us to assign both a causal weight and a direction of influence to each feature.

By combining SHAP values with gradient information, we can determine the causal impact and direction of each feature. The top-ranked causal features are then selected for further intervention or downstream analysis.

#### Optimization objectives and training strategy for CauFinder

Given the building modules described above, the learning of CauFinder can be formulated as minimizing the following overall loss *L*:

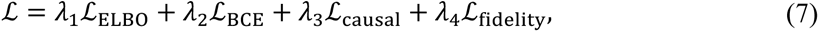

where *L*_ELBO_ is the negative evidence lower bound (ELBO) loss term that encourages the latent features *z* to stay in the data manifold; *L*_BCE_ ensures accurate prediction of the state or phenotype *y*, *L*_causal_ aims to maximize the causal information flow from *z*^*c*^ to *y*; *L*_fidelity_ ensures that *z*^*c*^ captures the main information about *y*; *λ*_1_, *λ*_2_, *λ*_3_ and *λ*_4_ are designed to balance these loss terms. To effectively balance these different loss components, we employ a staged training strategy that focuses on different losses at various stages (see Supplementary Note 7). Thus, our method is able to both theoretically and computationally identify the partial variables causally affecting the phenotypes/transitions among all observed variables.

The model was trained with the Adam optimizer^68^, with a mini-batch size of 128. The CauFinder architecture was implemented using PyTorch (v2.1.2)^69^ and run using a GPU running CUDA (v11.8). Please see the Supplementary Note 8 for more model architecture details.

### Identification of causal master regulators based on nonlinear network control

After determining the causal features and their respective causal weights, our next objective is to identify the master regulators that influence state or phenotype transitions. Given the complexity of biological systems, we incorporate prior network information to provide additional context and structural insights. To achieve this, we employ the classical network control algorithm, the Minimum Feedback Vertex Set (MFVS) method^25,26^, and integrate it with our causal weights. This approach helps us pinpoint key causal regulatory nodes within the state or phenotype transition network.

#### Prior gene interaction network construction

In this subsection, we construct a gene regulatory network by integrating multiple data sources to establish causal regulatory relations between genes. Following CEFCON, we use the comprehensive gene interaction network from NicheNet^70^ as our prior network. NicheNet includes ligand-receptor, intracellular signaling, and gene regulatory interactions from over 50 public mouse and human data sources. We focus on gene interactions within individual cells, removing ligand-receptor interactions.

We treat the unweighted network as directed by considering undirected edges as bidirectional. The original NicheNet network uses human gene symbols. For the mouse network, we map gene symbols using one-to-one orthologs from ENSEMBL, excluding ambiguous genes. This results in a human network with 25,332 genes and 5,290,993 edges, and a mouse network with 18,579 genes and 5,029,532 edges.

#### Minimum feedback vertex set method for identifying driver genes

The feedback vertex set (FVS)^25,26^ is a classical network control approach used for identifying driver genes with nonlinear characteristics, formulated as follows:

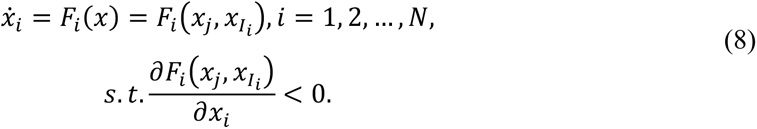

Here, *I_i_* represents the set of predecessor nodes of gene node *v_i_* (i.e., genes regulating *v*_i_), and the constraint condition ensures a decay property.

According to the method proposed by Zañudo et al.^71^, achieving control over all source nodes, which are nodes characterized by having an in-degree of zero, along with a strategically chosen set of feedback nodes, is sufficient to steer the system towards any of its potential attractors, representing different cell states. In the context of graph theory, the Feedback Vertex Set (FVS) problem is concerned with identifying a specific subset of nodes. The removal of these nodes from the graph ensures the elimination of all feedback cycles, effectively breaking any loops that may exist. This method is particularly suitable for the analysis and modeling of gene regulatory networks (GRNs). These networks are complex and intricate, often containing a multitude of both positive and negative feedback loops. Such feedback loops are integral to the functioning of natural biological circuits, playing a critical role in the regulation and maintenance of various cellular processes and states.

Our goal is to identify the minimum feedback vertex set while simultaneously maximizing the causal weights of the features we select. This means that we aim to pinpoint the smallest or least costly group of control nodes that also hold substantial significance in terms of causal relations. The problem can be formalized as the following 0-1 integer linear programming (ILP) optimization problem:

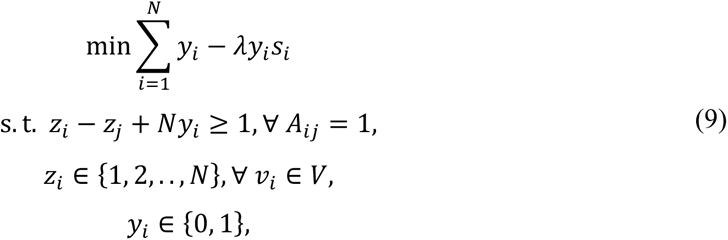

where *A* represents the adjacency matrix of the constructed network graph, where each entry *A_ij_* signifies the presence or absence of an edge between nodes. The variables *z_i_* and *z_j_* serve as auxiliary variables corresponding to nodes *v_i_* and *v_j_*, respectively. The binary variable *y_i_* is the decision variable, where *y_i_* = 1 indicates that node *v_i_* is included in the optimal feedback vertex set, and *y_i_* = 0 otherwise. The goal is to minimize the number of nodes in the feedback vertex set while accounting for their causal importance, as represented by *s_i_*.

Equation (9) describes a problem that falls into the category of NP-hard problems, meaning that it is unlikely to be solved efficiently using polynomial-time algorithms. In this study, we initially implement the graph reduction strategy as proposed by Lin and Jou to simplify the problem. Following this reduction, we employ the Gurobi optimizer (https://www.gurobi.com/) to solve the resulting ILP problem on the simplified graph.

By employing this approach, we are able to more precisely identify master regulators that play a critical role in orchestrating state or phenotype transitions. This methodology provides a robust theoretical foundation for designing targeted interventions, allowing us to effectively manipulate biological networks to achieve specific and desired state transitions.

#### Counterfactual generation for causal state transition

Counterfactual reasoning is a fundamental approach that helps individuals understand the consequences of their actions and the rules governing the world^72^. After identifying the causal master regulators *x*^*c*^and the causal regulatory pathway *y* = *f*^*c*^(*x*^*c*^), the challenge is how to control these genes to achieve desired state transitions. Here, *f*^*c*^represents the causal relation function from the causal master regulators *x*^*c*^to the final phenotype *y* . Specifically, the pathway is from *x*^*c*^ to *z*^*c*^ and then to *y*, where *z*^*c*^ is the causal latent variable extracted from *x*^*c*^. We formulate this problem as follows:

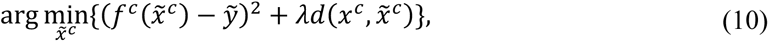

where *x*^*c*^ represents the causal variables of the target sample, *x̃*^*c*^ is the result after counterfactual or intervention, *ỹ* represents the desired outcome (state 0 or state 1, *ỹ* ∈ {0, 1}), and *λ* is used to balance the two distances. The term *d*(*x*^*c*^, *x̃*^*c*^) denotes the distance between the original instance *x*^*c*^ and the intervened instance *x̃*^*c*^.

This equation serves as the objective function for a heuristic counterfactual generation algorithm, aiming to balance the following two aspects. First, it seeks to make the intervened state *f*^*c*^(*x̃*^*c*^) as close as possible to the desired outcome *ỹ*. Second, it minimizes the distance between the original instance and the counterfactual instance, ensuring that the change from the original instance *x*^*c*^ to the intervened instance *x̃*^*c*^ is as small as possible.

By using this method, we aim to achieve significant semantic changes (state transitions) while making minimal perturbations to the basic elements of the causal variables. This approach allows for precise control of biological systems, enabling us to achieve the desired state transitions effectively.

#### Path controllability score

To evaluate the ease of controlling each sample during state transitions along a path within a fixed number of iterations *T*, we define the "path controllability score”. This metric is based on two factors: the change in master regulators (i.e., the amount of change in the key causal features of the sample during the transition process over *T* iterations) and the state change (i.e., the change in the sample’s state, such as the state probability change over *T* iterations). Path controllability is directly proportional to the state change and inversely proportional to the change in master regulators. This implies that if a sample achieves significant state change with smaller deviations in a fixed number of iterations, it is considered highly controllable. Based on these factors, we can define a simple formula for calculating the path controllability score:

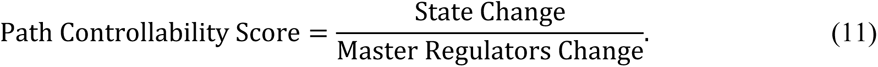

Here, the state change is defined as the difference between the state at iteration *T* and the initial state, and the master regulators change is the cumulative change in the master regulators over *T* iterations. This metric allows us to quantify the difficulty of controlling each sample within a fixed number of iterations, providing a useful measure for evaluating the effectiveness of our intervention strategies and the responsiveness of different samples to the applied controls.

### Datasets and data preprocessing

The human and mouse lung adenosquamous carcinoma (LUAS) sequencing datasets used in this study were sourced from the original article^59,60^. These datasets provide comprehensive sequencing data for 93 human LUAS samples and for mouse samples at various stages post-Ad-Cre administration: atypical adenomatous hyperplasia at 4 weeks, LUAD at 6 and 7 weeks, and LUSC at 8, 9, and 10 weeks. To facilitate our analysis, we applied a logarithmic transformation to the datasets. To facilitate our analysis, we applied a logarithmic transformation to the datasets.

For single-cell sequencing data, the data from previous study^62^ were utilized in our research, including scRNA-seq data with the Watermelon system, as well as processed cell information provided in the form of a meta-data matrix, which included critical information such as sequencing time, majority fate, and clone size. We preprocessed the data using Scanpy (v1.9.3) according to the methods provided in the original study. For each cell, we quantified the number of expressed genes and the proportion of transcripts from mitochondrial-encoded genes. Cells with fewer than 1,000 or more than 4,200 detected genes or a mitochondrial fraction greater than 0.1 were excluded from further analysis. Finally, the expression matrix was filtered to remove genes detected in fewer than three cells. This preprocessing resulted in a dataset containing 56,419 Watermelon-PC9 cells for the main analysis and an additional 16,477 cells from Watermelon models of EGFR-driven lung cancer (PC9) as an independent supplement, using 0.5 as cutoff for Minimum of dispersion leading to the identification of 1,296 highly variable genes. The preprocessed single-cell data were classified and labeled according to the provided cell information, resulting in several subsets used as input for CauFinder. Time-based groupings were classified according to known time labels, and cycling persister cells and non-cycling counterparts similarly classified based on given labels. For classifications not provided, such as cycling persister cells appearing on day 0, clonal barcodes were used to trace clonal lineages, enabling the tracking of cells’ clonal origin and their proliferative and transcriptional states.

## Data availability

All the datasets analyzed in this study are publicly available. The prior gene interaction network was derived from NicheNet^70^, which can be downloaded from https://github.com/saeyslab/nichenetr. Human and mouse embryonic stem cell single-cell RNA-seq datasets are available in the Gene Expression Omnibus (GEO) under accession codes GSE75748 (https://www.ncbi.nlm.nih.gov/geo/query/acc.cgi?acc=GSE75748) and GSE81682 (https://www.ncbi.nlm.nih.gov/geo/query/acc.cgi?acc=GSE81682), respectively. TCGA datasets used in this study can be downloaded from the UCSC Xena browser at https://xenabrowser.net/datapages/. Human Astrocytoma datasets are available in the NODE database under project accession number OEP001032 (https://www.biosino.org/node/sample/detail/OES032241). Mouse Astrocytoma datasets can be found in the NODE database under project accession number OEP002019 (https://www.biosino.org/node/project/detail/OEP002019). Single cell RNA-seq data for Watermelon system cells used in both cycling persisters and drug resistance studies, are available under GEO accession code GSE150949 (https://www.ncbi.nlm.nih.gov/geo/query/acc.cgi?acc=GSE150949). The data on drug responses to EGFR signaling pathway targeted therapies in individual patients and bulk RNA sequencing are derived from two databases: Genomics of Drug Sensitivity in Cancer^73^ (GDSC) and Cancer Cell Line Encyclopedia^74^ (CCLE). The GDSC data can be accessed and downloaded from (https://www.cancerrxgene.org/downloads/drug_data), while the CCLE data are available from (https://sites.broadinstitute.org/ccle/datasets). The Colorectal Cancer Liver dataset is sourced from website^63^ (http://www.cancerdiversity.asia/scCRLM).

## Code availability

CauFinder is an open-source collaborative initiative available in the GitHub repository (https://github.com/ChengmingZhang-CAS/CauFinder-main).

## Acknowledgements

This work was supported by National Basic Research Program of China (No. 2022YFA1004800); Strategic Priority Research Program of the Chinese Academy of Sciences 418 (No. XDB38040400); National Natural Science Foundation of China (Nos. T2341007, T2350003, 31930022,12131020 and 62002329); Special Fund for Science and Technology Innovation Strategy of Guangdong Province (Nos. 2021B0909050004, 2021B0909060002); JST (Japan Science and Technology Agency) Moonshot R&D (No. JPMJMS2021); AMED under Grant Number JP23dm0307009; Institute of AI and Beyond of the University of Tokyo; International Research Center for Neurointelligence (WPI-IRCN) at the University of Tokyo Institutes for Advanced Study (UTIAS); JSPS KAKENHI Grant Number JP20H05921;Henan Province Natural Science Foundation (242300421401), and Open Research Fund of State Key Laboratory of Digital Medical Engineering (2024-K07). We thank Dr. Shijie Tang and Yiwen Yang for their constructive comments on data analysis.

## Ethics declarations

Not applicable

