## Supplementary Information for "Steering cell-state and phenotype transitions by causal disentanglement learning"

#### Supplementary Notes

##### Supplementary Note 1: Simulation dataset construction

The simulated dataset was constructed to explore the effects of both causal and spurious relationships among observed variables. In this simulation, we focus on three types of observed variables: $x^c$, $x^s$, and $y$. The variables $x^c$ have a direct causal effect on the outcome $y$, while the spurious variables $x^s$ are correlated with $y$ without having a direct causal effect. Additionally, the outcome variable $y$ is directly observed. The unobserved variable $u \sim N(\mu^u, \sigma^u)$ acts as a confounder, creating dependencies among the observed variables. The primitive variable $c \sim N(\mu^c, \sigma^c)$ directly influences the observed variables with a causal effect. The spurious variables $x^s$ are generated by the function $f_1(u)$. Although $x^s$ has no direct causal effect on the outcome variable $y$, it is correlated with $y$ through the shared dependence on $u$. The observed variables with a causal effect on $y$, denoted as $x^c$, are generated by combining the effects of $f_2(x^s)$ and $g_1(c)$. This means that some $x^c$ can be influenced by both partial spurious variables and the primitive causal variable, reflecting the complexity often observed in real-world data. The outcome variable $y$ is modeled as a composite of multiple effects. Specifically, $y$ is expressed as $\lambda g_2(x^c) + (1 - \lambda) f_3(u) + \gamma\epsilon$, where $\lambda$ controls the strength of the causal effect from $x^c$, $\gamma$ controls the noise scale, and $\epsilon$ is a non-Gaussian noise to ensure the directed causal relationship[1] from $x$ to $y$. In this construction, $\lambda$ plays a crucial role in balancing the contributions of $x^c$ and $u$ to $y$. When $\lambda$ is close to 1, the causal effect of $x^c$ dominates, whereas when $\lambda$ is close to 0, the effect of the unobserved variable $u$ is more prominent. The noise term $\gamma\epsilon$ adds randomness to $y$, simulating the inherent variability observed in real-world data.

In the linear scenario, $f_1, f_2, f_3, g_1, g_2$ are all functions where a random weight is multiplied by the input. In the nonlinear scenario, the input is further transformed using the hyperbolic tangent function. The sample size is set to 200, with 90 confounding features ($x^s$) and 10 causal features ($x^c$).

##### Supplementary Note 2: Benchmark methods and software programs

In our benchmark experiments, we employed several methods to evaluate the extraction of feature subsets with causal effects on the state variable $y$ from all observed variables. These methods can be categorized into three groups: causal inference methods, deep learning models, and machine learning models.

**Causal model:** we used the PC algorithm[2], which constructs a directed causal network based on the input variables, focusing only on those that have a causal effect on $y$ to ensure a relevant comparison.

**Deep learning models:** We used the variational autoencoder (VAE), implemented in PyTorch (v2.1.2)[3]. For the VAE model, we set the number of hidden neurons to 64 and the latent space variables to 10. Similar to CauFinder, we used the latent space to predict $y$ during training. We assigned weights to each feature using SHAP values (VAE_SHAP) or by calculating the gradient of each feature (VAE_Grad). This approach can be treated as an ablation study of CauFinder since it lacks the causal component.

**Machine learning models:** Implemented using Python's statistic machine learning libraries such as scikit-learn (v1.13)[4], we used T-test, random forest (RF), mutual information (MI), Pearson correlation coefficients (PCC), and Spearman correlation coefficients (SCC). For the Random Forest model, we set the number of trees to 50 and the maximum depth to 2. Causal features were selected based on computed feature importance.

Additionally, we used the following methods for comparisons with hESC and mHSC data benchmark evaluation:

**CellOracle**: CellOracle[5] utilizes single-cell RNA sequencing (scRNA-seq) data and a prior base gene regulatory network (GRN), or a combination of scRNA-seq and single-cell ATAC sequencing (scATAC-seq) data to compute the base GRN. Based on this base GRN, it constructs cell type-specific GRNs and employs various strategies to identify key transcription factors (TFs). These key TFs are then subjected to simulated perturbations to predict changes in cell states. In our comparisons, we followed the strategy outlined in CEFCON, using the default degree centrality of TFs in the GRN to measure their importance.

**CEFCON**: CEFCON[6] is a network framework for inferring gene regulatory relationships and characterizing their dynamic properties from the perspective of network control theory, aimed at identifying driver regulatory factors in cell fate decisions. CEFCON takes prior gene interaction networks and scRNA-seq data as inputs and combines network attention coefficients, minimum dominating set (MDS), and minimum feedback vertex set (MFVS) methods to obtain a final influence score for each gene. This score serves as the criterion for determining whether a gene is a driver of state transitions.

**WMDS.net**: WMDS.net[7] is a weighted minimum dominating set network model based on structural controllability theory. It identifies the minimum dominating set of driver nodes in a transcriptional co-expression network as critical drivers for network state transitions.

WMDS.net combines the node degree and the significance of differential co-expression of genes between two states to measure node controllability within the transcriptional network.

#### Supplementary Note 3: Evaluation criteria on benchmark datasets

For the evaluation of our models on benchmark datasets, we employed a variety of metrics to ensure comprehensive performance assessment. These metrics include:

**Accuracy (ACC):** The proportion of true results (both true positives and true negatives) among the total number of cases examined. It is calculated as $\frac{TP+TN}{TP+TN+FP+FN}$, where TP is true positives, TN is true negatives, FP is false positives, and FN is false negatives.

**Area Under the Receiver Operating Characteristic Curve (AUC):** Measures the ability of the model to distinguish between classes. A higher AUC value indicates better model performance. The AUC ranges from 0 to 1, where 1 represents a perfect model and 0.5 represents a model with no discriminative ability.

**Recall (Sensitivity or True Positive Rate):** The proportion of actual positives correctly identified by the model. It is calculated as $\frac{TP}{TP+FN}$ . High recall indicates that the model is able to identify most of the positive cases (causal features).

**Specificity (True Negative Rate):** The proportion of actual negatives correctly identified by the model. It is calculated as $\frac{TN}{TN+FP}$. High specificity indicates that the model is able to identify most of the negative cases (spurious features).

**Precision (Positive Predictive Value):** The proportion of positive identifications that are actually correct. It is calculated as $\frac{TP}{TP+FP}$. High precision indicates that the model has a low false positive rate.

**F1 Score:** The harmonic mean of precision and recall, providing a balance between the two. It is calculated as $F1 = 2\frac{\text{Precision.Recall}}{\text{Precision+Recall}}$. This metric is particularly useful when the class distribution is imbalanced.

**Matthews Correlation Coefficient (MCC):** A correlation coefficient that takes into account true and false positives and negatives, providing a balanced measure even if the classes are of very different sizes. It is calculated as $MCC = \frac{TP.TN-FP.FN}{\sqrt{(TP+FP)(TP+FN)(TN+FP)(TN+FN)}}$. The MCC value ranges from -1 to +1, where +1 indicates a perfect prediction, 0 indicates no better than random prediction, and -1 indicates total disagreement between prediction and observation.

These metrics provide a comprehensive understanding of the model's performance, highlighting strengths and potential areas for improvement.

#### Supplementary Note 4: Choosing the number of causal drivers

Determining the appropriate threshold to identify the number of causal drivers is challenging. Typically, CauFinder assigns a causal weight to each feature, with higher values indicating a greater likelihood of being a causal driver.

For evaluating performance on simulated data, we explored two methods. The first method involves selecting the top 10 features as causal drivers. The second method involves ranking the features by their causal weights in descending order and then summing these weights cumulatively, selecting the top 30% of features based on this cumulative proportion. For clarity and alignment with the simulated data's construction, which includes 10 causal features, we opted to use the first method, selecting the top 10 features as causal drivers for all benchmark methods. In real-world scenarios, we combine prior knowledge and control theory to assist in this selection.

#### Supplementary Note 5: Derivation of the estimator of information flow

This section provides the detailed derivation of the estimator for the causal information flow $I(z^c \to y)$.

The causal information flow between the causal factors $z^c$ and the prediction $y$ can be calculated as:

$$I(z^c \to y) = \int_{z^c} P(z^c) \sum_y P\big(y|do(z^c)\big) \log \frac{P\big(y|do(z^c)\big)}{\int_{z^c} P(z^c) P\big(y|do(z^c)\big) dz^c} dz^c. \tag{S1}$$

Expanding this, we have:

$$\begin{aligned} I(z^c \to y) = & \int_{z^c} P(z^c) \left( \sum_y P\big(y|do(z^c)\big) \log P\big(y|do(z^c)\big) \right) dz^c \\ & - \sum_y \int_{z^c} P(z^c) P\big(y|do(z^c)\big) dz^c . \log \int_{z^c} P(z^c) P\big(y|do(z^c)\big) dz^c. \end{aligned} \tag{S2}$$

Next, we compute $P\big(y|do(z^c)\big)$, which can be efficiently estimated using Monte Carlo sampling. Specifically, we have:

$$P\big(y|do(z^c)\big) = \sum_x P(y|z^c,x)P(x) = \sum_x \int_{z^s} P(z^s|z^c,x)P(y|z^c,z^s)P(x)dz^s$$
$$\approx \frac{1}{N^x N^s}\sum_{k=1}^{N^x}\sum_{j=1}^{N^s} P\big(y|z^c,z^s_{kj}\big). \tag{S3}$$

where $k$ indexes the $N^x$ samples $x_k$ drawn from the dataset, and $j$ indexes the $N^s$samples for each $x_k$, i.e., $z^s_{kj}\sim P(z^s|z^c,x_k)$.Here, we approximate the true posterior distribution $P(z^s|z^c,x_k)$ using the variational distribution $q(z^s|z^c,x_k)$. Note that in Equation (S3), $x$, $z^c$, and $z^s$ do not necessarily belong to the same sample from the original dataset. Then, we have

$$\int_{z^c} P(z^c)P\big(y|do(z^c)\big)dz^c$$
$$= \int_{z^c}\sum_x \int_{z^s} P(x)P(z^c)P(z^s|z^c,x)P(y|z^c,z^s)dz^s\, dz^c$$
$$\approx \frac{1}{N^c N^x N^s}\sum_{i=1}^{N^c}\sum_{k=1}^{N^x}\sum_{j=1}^{N^s} P\big(y|z^c_i,z^s_{kj}\big). \tag{S4}$$

Similarly, $i$ indexes the $N^c$ samples from $z^c$'s marginal distribution, i.e., $z^c_i\sim P(z^c)$, $k$indexes the $N^x$ samples from $x$'s marginal distribution $P(x)$, and $j$ indexes the $N^s$ samples of $z^s$ for each pair $(z^c_i,x_k)$, i.e., $z^s_{ikj}\sim P(z^s|z^c_i,x_k)$. In practice, we approximate the true posterior distribution $P(z^s|z^c_i,x_k)$ using the variational distribution $q(z^s|z^c_i,x_k)$. Combining these, we get:

$$I(z^c \to y)$$
$$= \frac{1}{N^c}\sum_{i=1}^{N^c}\sum_y\left(\frac{1}{N^x N^s}\sum_{k=1}^{N^x}\sum_{j=1}^{N^s}P\big(y|z^c_i,z^s_{kj}\big)\right)\log\left(\frac{1}{N^x N^s}\sum_{k=1}^{N^x}\sum_{j=1}^{N^s}P\big(y|z^c_i,z^s_{kj}\big)\right)$$
$$-\sum_y\left(\frac{1}{N^c N^x N^s}\sum_{i=1}^{N^c}\sum_{k=1}^{N^x}\sum_{j=1}^{N^s}P\big(y|z^c_i,z^s_{kj}\big)\right).\log\left(\frac{1}{N^c N^x N^s}\sum_{i=1}^{N^c}\sum_{k=1}^{N^x}\sum_{j=1}^{N^s}P\big(y|z^c_i,z^s_{kj}\big)\right)$$
$$= \frac{1}{N^c N^x N^s}\left[\sum_{i=1}^{N^c}\sum_y\left(\sum_{k=1}^{N^x}\sum_{j=1}^{N^s}P\big(y|z^c_i,z^s_{kj}\big)\right).\log\left(\frac{1}{N^x N^s}\sum_{k=1}^{N^x}\sum_{j=1}^{N^s}P\big(y|z^c_i,z^s_{kj}\big)\right)\right.$$
$$\left.-\sum_y\left(\sum_{i=1}^{N^c}\sum_{k=1}^{N^x}\sum_{j=1}^{N^s}P\big(y|z^c_i,z^s_{kj}\big)\right).\log\left(\frac{1}{N^c N^x N^s}\sum_{i=1}^{N^c}\sum_{k=1}^{N^x}\sum_{j=1}^{N^s}P\big(y|z^c_i,z^s_{kj}\big)\right)\right]. \tag{S5}$$

#### Supplementary Note 6: Impact of unobserved variables on information flow

In our current framework, we have considered scenarios where all variables $x$ are observed. However, a natural question arises: what happens if some variables are unobserved? This situation is common in real-world data, as it is often impossible to observe every variable. To address this, we refined our division of features in both the latent and original spaces, by introducing $u$ to represent unobserved variables (Supplementary Fig. 2). We further split the original $z^c$ into $z_{xu}^c$, representing the causal latent space shared by $x$ and $u$, and $z_x^c$, specific to $x$. Additionally, $z_u^c$ denotes the causal latent space unique to $u$. Consistent with our previous notation, $z^c = \{z_{xu}^c, z_x^u\}$. In this context, the causal latent space comprises both $z^c$ and $z_u^c$. A similar approach is applied to defining spurious latent spaces.

The core of our exploration focuses on the impact of unobserved variables $u$ on the causal information flow $I(z^c \to y)$. This inquiry centers on $u$'s effect on $P(y|do(z^c))$, the conditional distribution given the intervention on $z^c$ using do-calculus. From a probabilistic graphical perspective, compared to the case where $uu$ is absent, four additional paths are introduced: $u \to z_{xu}^c; u \to z_u^c;\ u \to z_{xu}^s; u \to z_u^s$. Although there are $2^4 - 1$ possible combinations of the presence and absence of these latent variables, the primary distinction lies in whether $z_u^c$ is present or absent. Therefore, we focus our discussion on these two scenarios: when $z_u^c$ is present and when $z_u^c$ is absent. We consider two scenarios: when $z_u^c$ is present and when $z_u^c$ is absent.

(1) When $z_u^c$ is present, the causal structure model can be represented as: $x, u \to z_{xu}^c; x, u \to z_{xu}^s; x \to z_x^c; x \to z_x^s; u \to z_u^c;\ u \to z_u^s; z_u^c, z^c, z^s \to y$. In this case, the conditional distribution $P(y|do(z^c))$ is derived as follows:

$$
\begin{aligned}
P\big(y|do(z^c)\big) &= \sum_{x,u} P(y|z^c,x,u)P(x,u) \\
&= \sum_{x,u} \int_{z^s}\int_{z_u^s}\int_{z_u^c} P(z^s|x,u)P(z_u^c|u)P(z_u^s|u)P(y|z^c,z^s,z_u^c)P(x,u)\, dz^s dz_u^s dz_u^c \\
&= \sum_{x,u} \int_{z^s}\int_{z_u^c} P(z^s|x)P(z_u^c|u)P(y|z^c,z^s,z_u^c)P(u|x)P(x)\, dz^s dz_u^c \\
&= \sum_{x} \int_{z^s} P(z^s|x)P(x)\left(\sum_{u}\int_{z_u^c} P(y|z^c,z^s,z_u^c)\, P(u|x)dz_u^c\right) dz^s \\
&= \sum_{x} \int_{z^s} P(z^s|z^c,x)\tilde{P}(y|z^c,z^s)P(x)dz^s \\
&\approx \frac{1}{N^x N^s}\sum_{k=1}^{N^x}\sum_{j=1}^{N^s} \tilde{P}\big(y|z^c,z_{kj}^s\big).
\end{aligned}
\tag{S6}
$$

Here, $\tilde{P}(y|z^c,z^s) = \sum_u \int_{z_u^c} P(y|z^c,z^s,z_u^c)\, P(u|x)dz_u^c$ represents the average effect of $z_u^c$ on $y$. We assume that $P(z^s|x,u) = P(z^s|x)$, leveraging the conditional independence assumption. This means that $x$ contains all the necessary information about $z^s$, and thus the influence of $u$ on $z^s$ can be ignored. Since $u$ is unobserved, this probability is not directly computable, making the calculation of $I(z^c \to y)$ potentially inaccurate in this scenario.

(2) When $z_u^c$ is absent, the causal structure model can be represented as: $x,u \to z_{xu}^c; x,u \to z_{xu}^s; x \to z_x^c; x \to z_x^s; u \to z_u^c;\ u \to z_u^s; z_u^c, z^c, z^s \to y$. In this case, the conditional distribution $P\big(y|do(z^c)\big)$ is derived as follows:

$$
\begin{aligned}
P\big(y|do(z^c)\big) &= \sum_{x,u} P(y|z^c,x,u)P(x,u) \\
&= \sum_{x,u} \int_{z^s}\int_{z_u^s} P(z^s|x,u)P(z_u^s|u)P(y|z^c,z^s)P(x,u)\, dz^s dz_u^s \\
&= \sum_{x,u} \int_{z^s} P(z^s|x,u)P(y|z^c,z^s)P(x,u)dz^s \\
&= \sum_{x} \int_{z^s} P(z^s|z^c,x)P(y|z^c,z^s)P(x)\left(\sum_{u} P(u|x)\right) dz^s \\
&= \sum_{x} \int_{z^s} P(z^s|z^c,x)P(y|z^c,z^s)P(x)dz^s \\
&\approx \frac{1}{N^x N^s}\sum_{k=1}^{N^x}\sum_{j=1}^{N^s} P\big(y|z^c,z_{kj}^s\big).
\end{aligned}
\tag{S7}
$$

This demonstrates that, in this scenario, the presence of unobserved variables $u$ does not affect $P\big(y|do(z^c)\big)$, and therefore, does not impact the defined causal information flow $I(z^c \rightarrow y)$.

In conclusion, our analysis confirms that the definition of $I(z^c \rightarrow y)$ holds accurate when $x$ captures all essential causal information concerning $y$. This condition is typically met in high-dimensional feature spaces, such as scRNA-seq data, where a comprehensive encapsulation of causal information relative to $y$ is achievable. This ensures the reliability of causal inferences in complex biological datasets where not all variables may be observable.

#### Supplementary Note 7: Staged training strategy

To effectively balance these different loss components, we employ a staged training strategy that focuses on different losses at various stages.

In the initial stage, approximately the first 10% of the epochs, the primary focus is on minimizing the ELBO loss $\mathcal{L}_{\text{ELBO}}$, specifically emphasizing the reconstruction losses for the causal and spurious VAEs ($\mathcal{L}_{\text{rec1}}$ and $\mathcal{L}_{\text{rec2}}$). This stage aims to embed the latent features $z$ within the data manifold and ensure accurate data reconstruction. The loss for $\mathcal{L}_{\text{BCE}}$, $\mathcal{L}_{\text{causal}}$ and $\mathcal{L}_{\text{fidelity}}$ are not considered in this phase.

From approximately 10% to 40% of the epochs, the emphasis within the ELBO loss expands to include the KL divergence loss. This transition helps in regularizing the latent space by enforcing a prior distribution over the latent variables, which is crucial for a well-behaved latent space.

From approximately 40% to 70% of the epochs, the focus extends to include the binary cross-entropy (BCE) loss $\mathcal{L}_{\text{BCE}}$. This phase aims to enhance the accuracy of phenotype or state predictions by directly optimizing for the prediction task.

In the final stage, constituting approximately the final 30% of the epochs, the focus extends to include minimizing the causal-related losses, specifically the causal loss $\mathcal{L}_{\text{causal}}$ and the fidelity loss $\mathcal{L}_{\text{fidelity}}$. This phase ensures that the causal relationships are accurately captured and the representation of causal features is refined.

This staged approach ensures a balanced and comprehensive optimization process, enabling CauFinder to effectively distinguish and manipulate causal factors.

#### Supplementary Note 8: CauFinder architecture

CauFinder employs a Dual Variational Autoencoder (DVAE) architecture, implemented using the PyTorch (version 1.13.0)[3] and Scanpy (version 1.9.1)[8] Python libraries. This architecture integrates several key components to facilitate causal modeling and network control, comprising a feature selection layer, an encoder, a decoder, and a classifier for binary cross-entropy (BCE) loss.

The feature selection layer is the first layer applied to the input data $=x$. It splits the input features into causal and spurious components using initial weights and a threshold. An optional attention mechanism, implemented through an attention network with two linear layers and LeakyReLU activations, refines the feature selection process.

The encoder projects the input vector to 128 dimensions, followed by BatchNorm1d batch normalization, ReLU non-linear activation, and Dropout regularization with a dropout rate of 0.1. It consists of two modules: one projects causal features $x_1$ into a latent space $z^c$ with 2 dimensions (default), and the other projects spurious features $x_2$ into a latent space $z^s$ with 8 dimensions. Both modules utilize multiple layers, batch normalization, ReLU activation, and the reparameterization trick for sampling.

The decoder mirrors the encoder architecture. It first applies a fully connected layer and ReLU activation to project the latent representation back to 128 dimensions, then projects this 128-dimensional layer into outputs matching the input vector size. The decoder also comprises two modules: one reconstructs $x_{rec1}$ from $z^c$, and the other reconstructs $x_{rec2}$ from $z^s$. The decoder employs a dropout rate of 0.0.

The classifier for binary cross-entropy (BCE) loss predicts the phenotype or state $y$. This classifier uses the latent representations $z^c$ and $z^s$ to perform binary classification, enhancing the accuracy of phenotype or state predictions. The classifier applies a fully connected layer to the concatenated latent vectors and outputs the predicted probabilities.

This DVAE architecture, combined with the feature selection layer and classifier, ensures that CauFinder effectively captures and manipulates causal and spurious factors, enabling precise guidance of phenotype transitions.

### Supplementary Figures

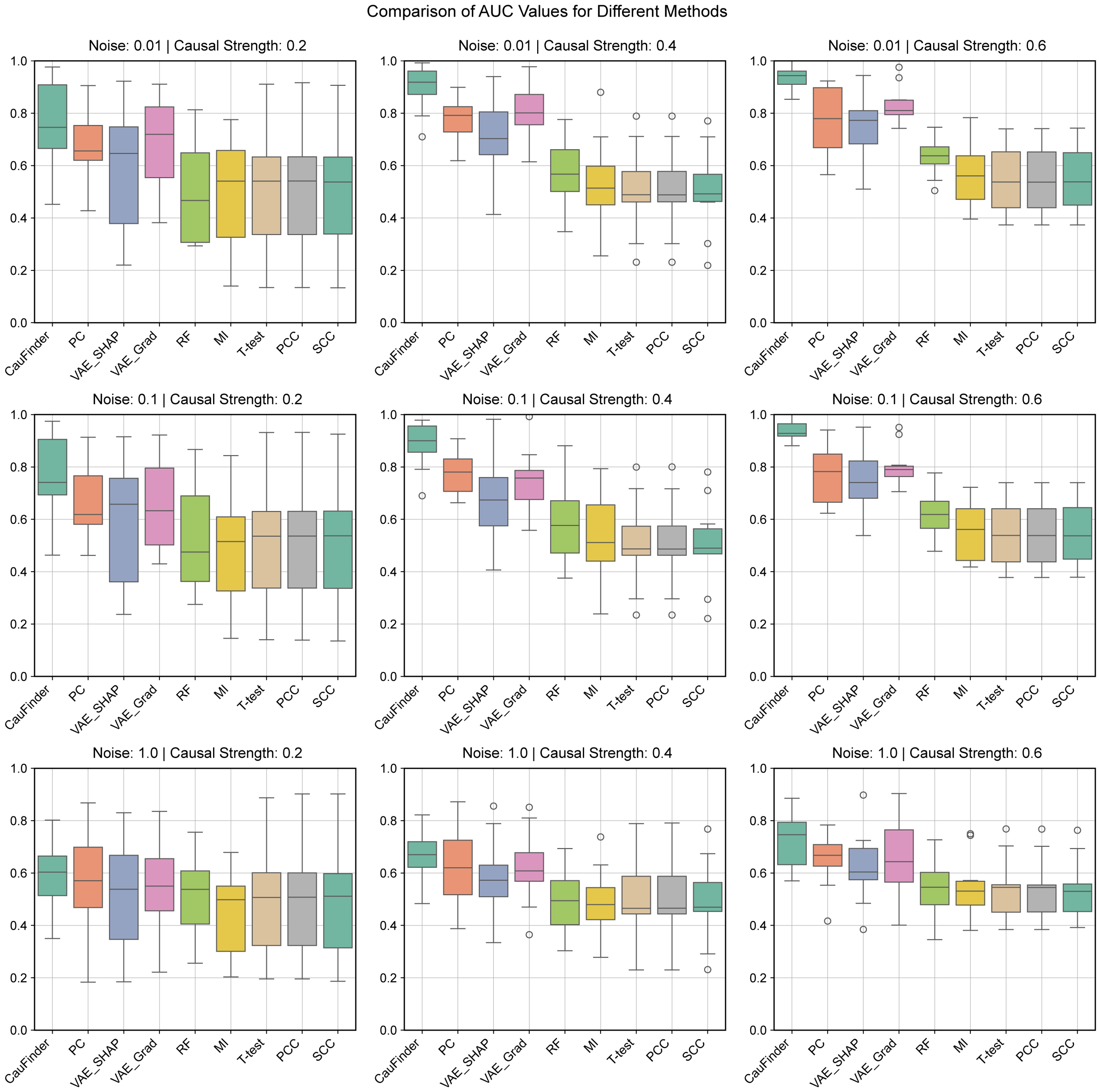

**Supplementary Fig. 1: Comparison of AUC values for different methods across various causal strengths and noise levels based on linear simulated data.** The nine-panel box plot grid displays the AUC (Area Under the Curve) values for our model and other competing models under different simulated conditions. Each panel represents a specific combination of causal strength (0.2, 0.4, 0.6) and noise level (0.01, 0.1, 1.0). The methods compared include PC, VAE-SHAP, VAE-Grad, RF, MI, T-test, PCC, and SCC. The y-axis in each box plot represents the AUC values, illustrating the performance variability and robustness of each method under varying causal strengths and noise levels, based on linear simulated data.

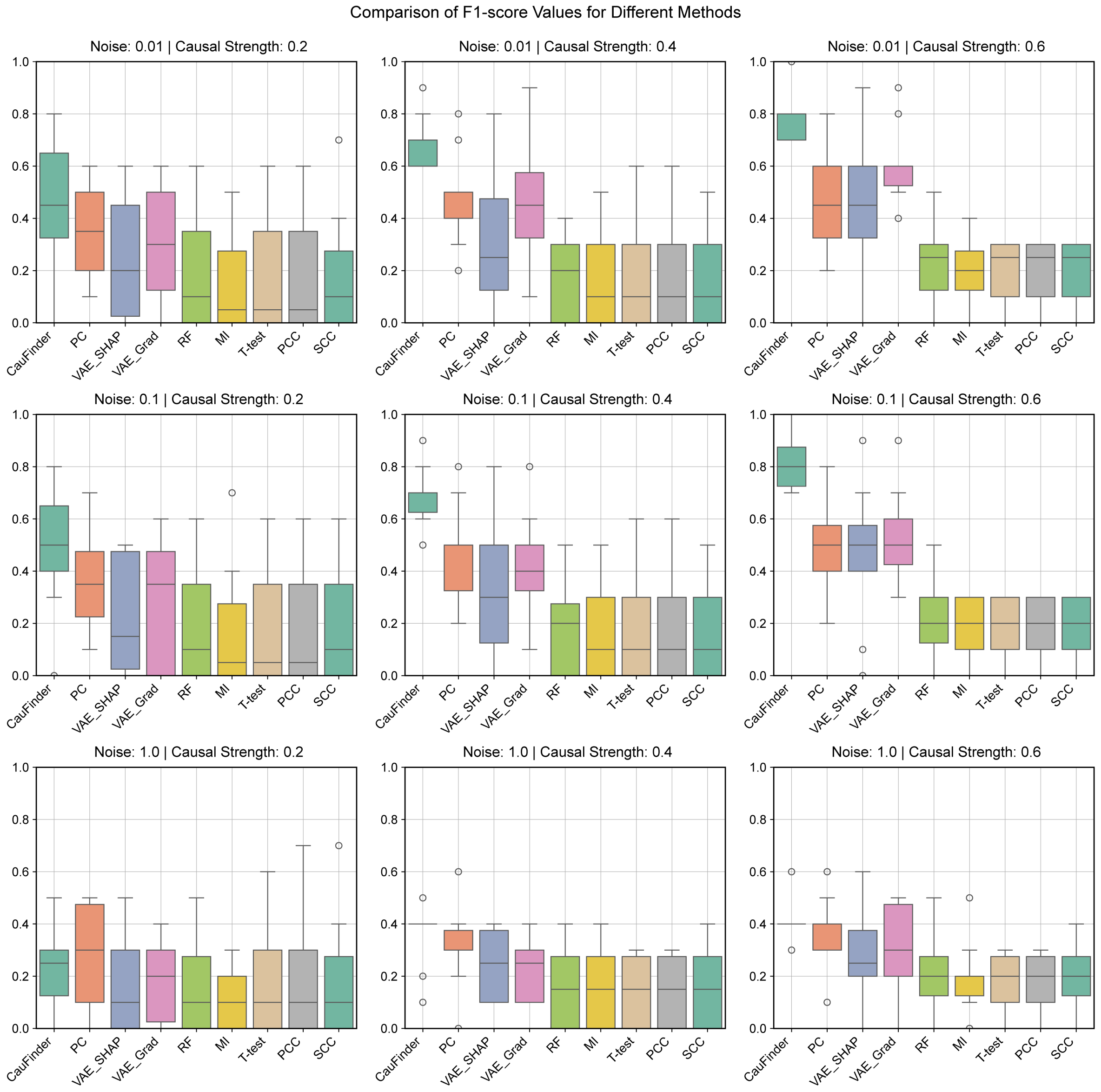

**Supplementary Fig. 2: Comparison of F1-score values for different methods across various causal strengths and noise levels based on linear simulated data.** The nine-panel box plot grid displays the F1-score values for our model and other competing models under different simulated conditions. Each panel represents a specific combination of causal strength (0.2, 0.4, 0.6) and noise level (0.01, 0.1, 1.0). The methods compared include PC, VAE-SHAP, VAE-Grad, RF, MI, T-test, PCC, and SCC. The y-axis in each box plot represents the F1-scores, illustrating the performance variability and robustness of each method under varying causal strengths and noise levels, based on linear simulated data.

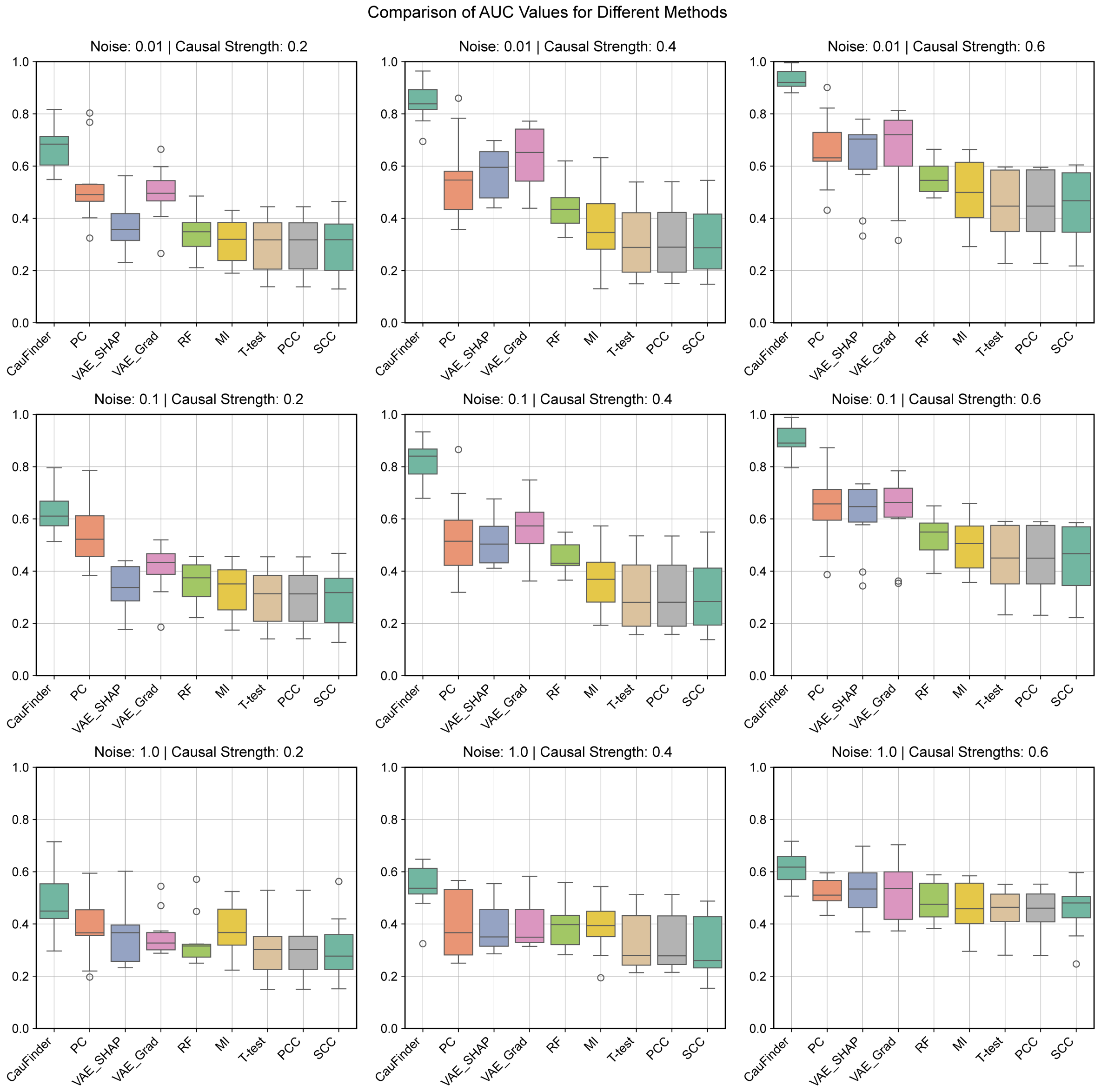

**Supplementary Fig. 3: Comparison of AUC values for different methods across various causal strengths and noise levels based on nonlinear simulated data.** The nine-panel box plot grid displays the AUC (Area Under the Curve) values for our model and other competing models under different simulated conditions. Each panel represents a specific combination of causal strength (0.2, 0.4, 0.6) and noise level (0.01, 0.1, 1.0). The methods compared include PC, VAE-SHAP, VAE-Grad, RF, MI, T-test, PCC, and SCC. The y-axis in each box plot represents the AUC values, illustrating the performance variability and robustness of each method under varying causal strengths and noise levels, based on nonlinear simulated data.

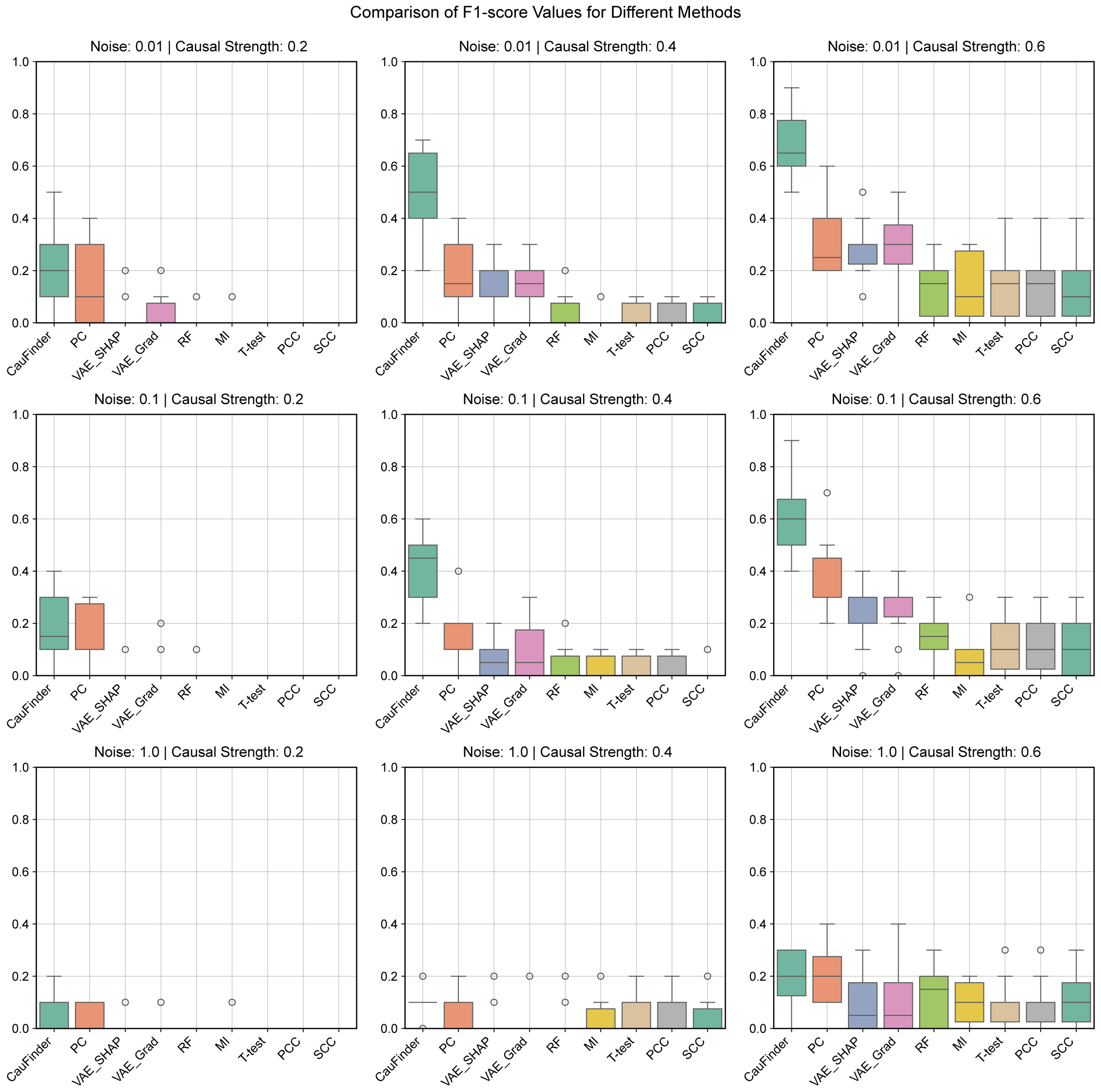

**Supplementary Fig. 4: Comparison of F1-score values for different methods across various causal strengths and noise levels based on nonlinear simulated data.** The nine-panel box plot grid displays the F1-score values for our model and other competing models under different simulated conditions. Each panel represents a specific combination of causal strength (0.2, 0.4, 0.6) and noise level (0.01, 0.1, 1.0). The methods compared include PC, VAE-SHAP, VAE-Grad, RF, MI, T-test, PCC, and SCC. The y-axis in each box plot represents the F1-scores, illustrating the performance variability and robustness of each method under varying causal strengths and noise levels, based on nonlinear simulated data.

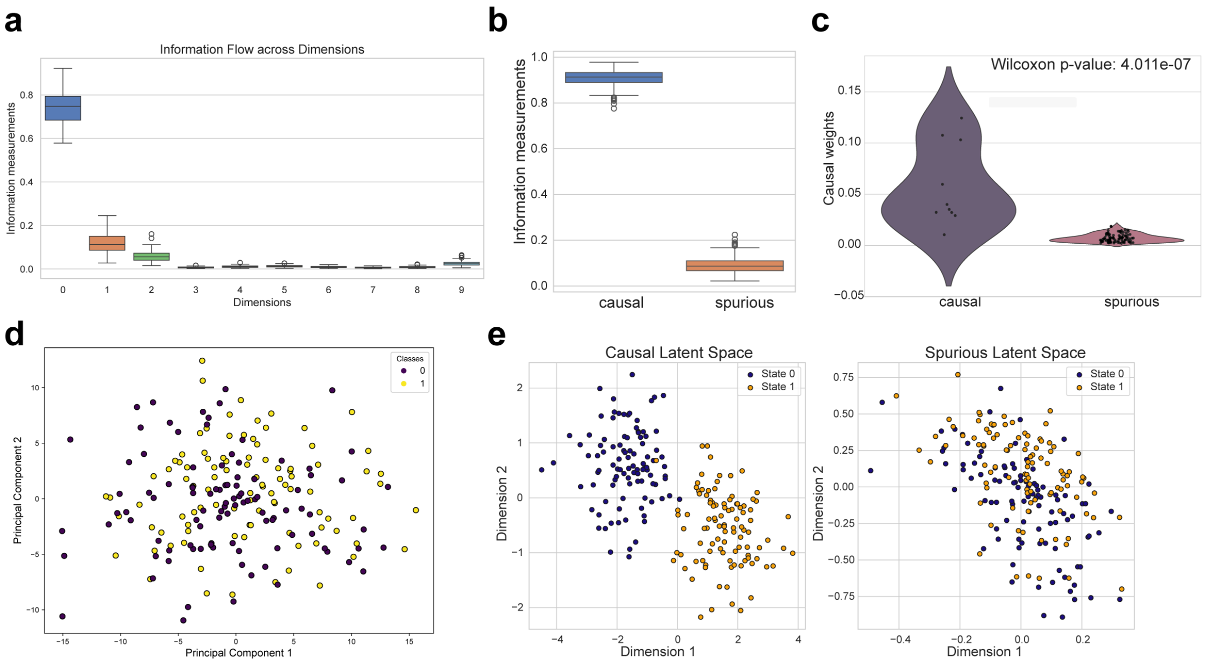

**Supplementary Fig. 5: Causal decoupling performance of CauFinder on simulated data. a,** Causal information flow across different dimensions of the latent space. The x-axis represents the dimensions, and the y-axis represents the causal information flow values. Each boxplot point represents a sample. **b,** Boxplot illustrating the distribution of information measurements for the causal dimensions (left) and spurious dimensions (right) in the model's latent space. The y-axis represents the information measurements. **c,** Violin plot displays the distribution of feature weights assigned by the model to causal features (left) and spurious features (right), with violin width indicating feature density. **d,** Principal component analysis plot of the simulated data. The x-axis and y-axis represent the first two principal components, respectively. Colours indicate the two classes (Class 0 and Class 1). **e**, Scatter plots compare the sample distributions in the model's latent space based on causal dimensions (left) and spurious dimensions (right). Each point represents a sample, colored by class (Class 0 in blue and Class 1 in orange).

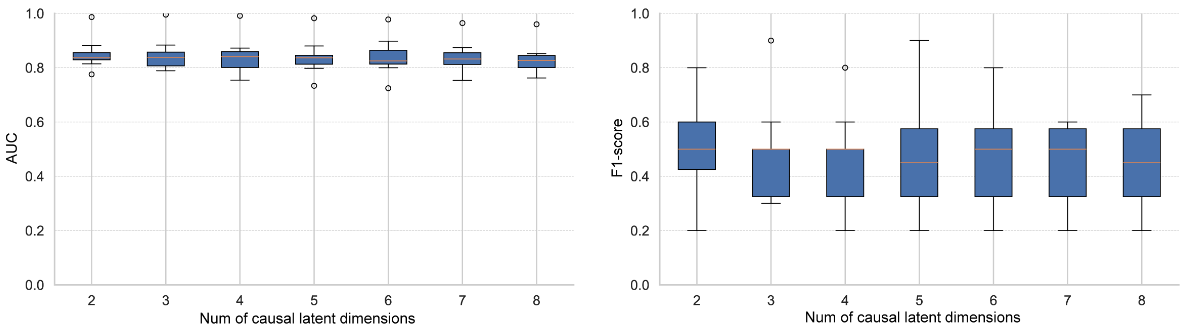

**Supplementary Fig. 6: The effect of different numbers of causal latent spaces on model performance metrics.** In (a), the x-axis represents the number of causal latent spaces (n_causal), and the y-axis represents the AUC value. In (b), the x-axis represents the number of causal latent spaces (n_causal), and the y-axis represents the F1 score.

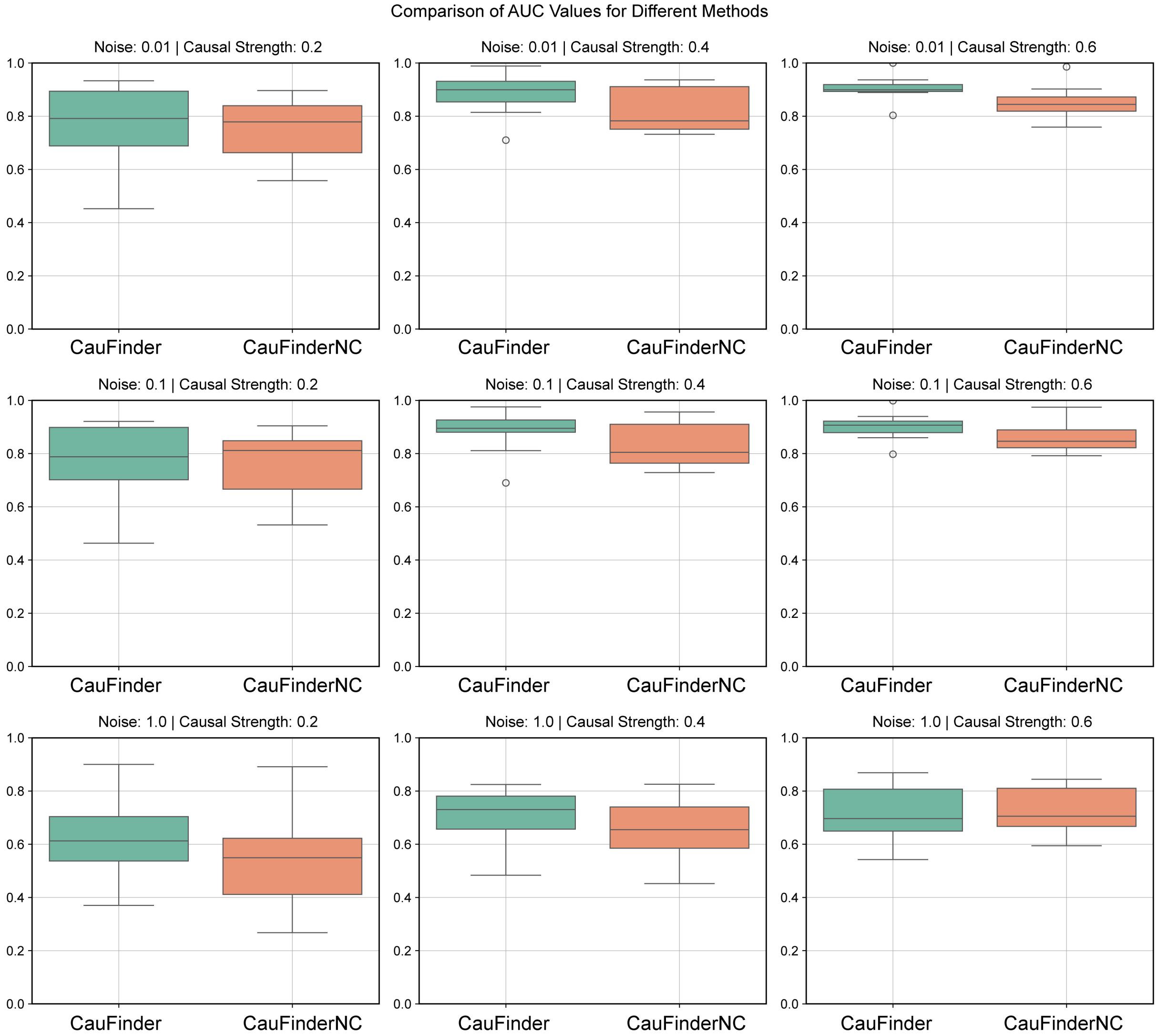

**Supplementary Fig. 7: Results of ablation experiment with and without causal weights.** This figure shows the performance comparison between the algorithm with causal weights (CauFinder) and without causal weights (CauFinderNC).

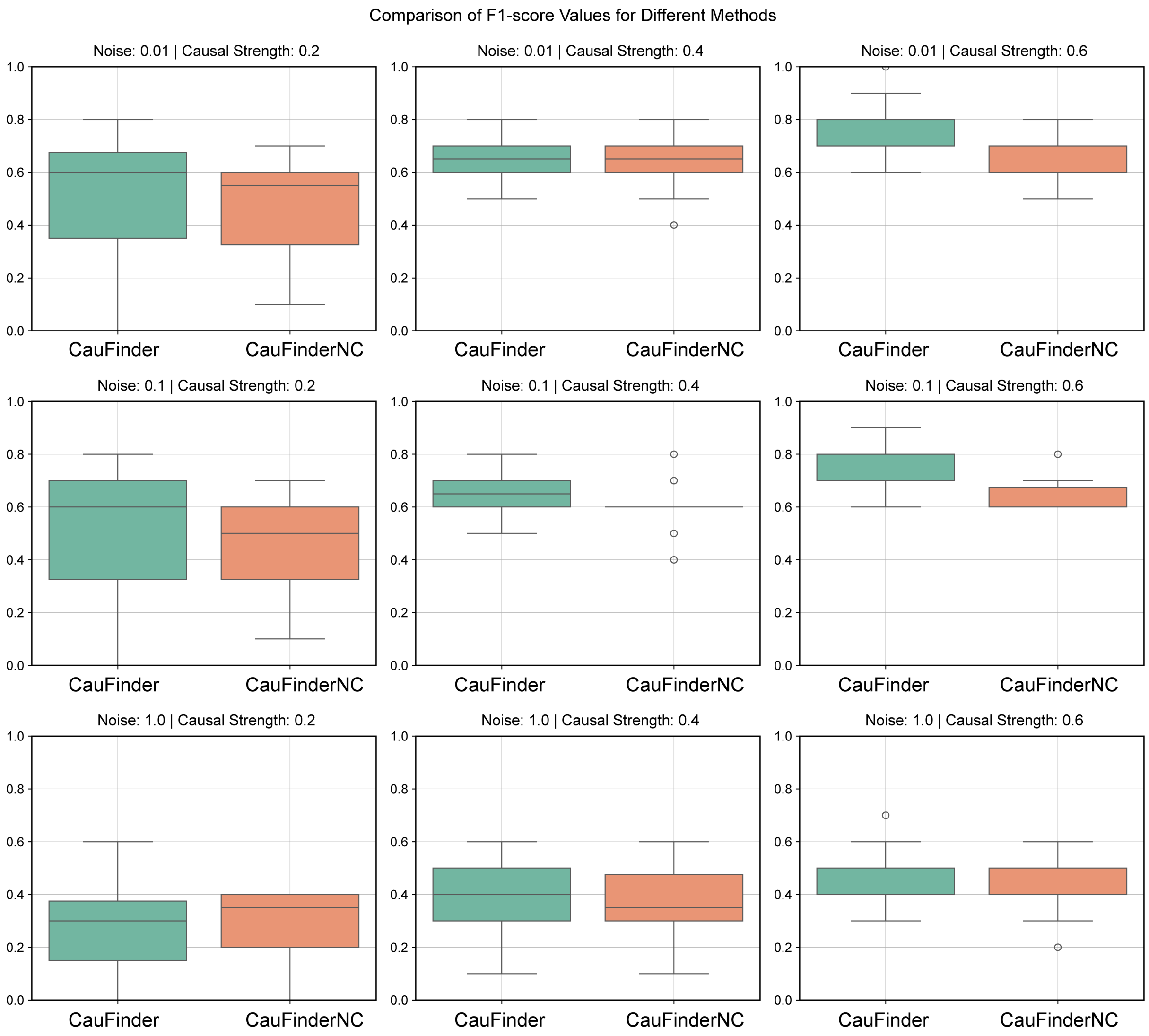

**Supplementary Fig. 8: Results of ablation experiment with and without causal weights.** This figure shows the performance comparison between the algorithm with causal weights (CauFinder) and without causal weights (CauFinderNC).

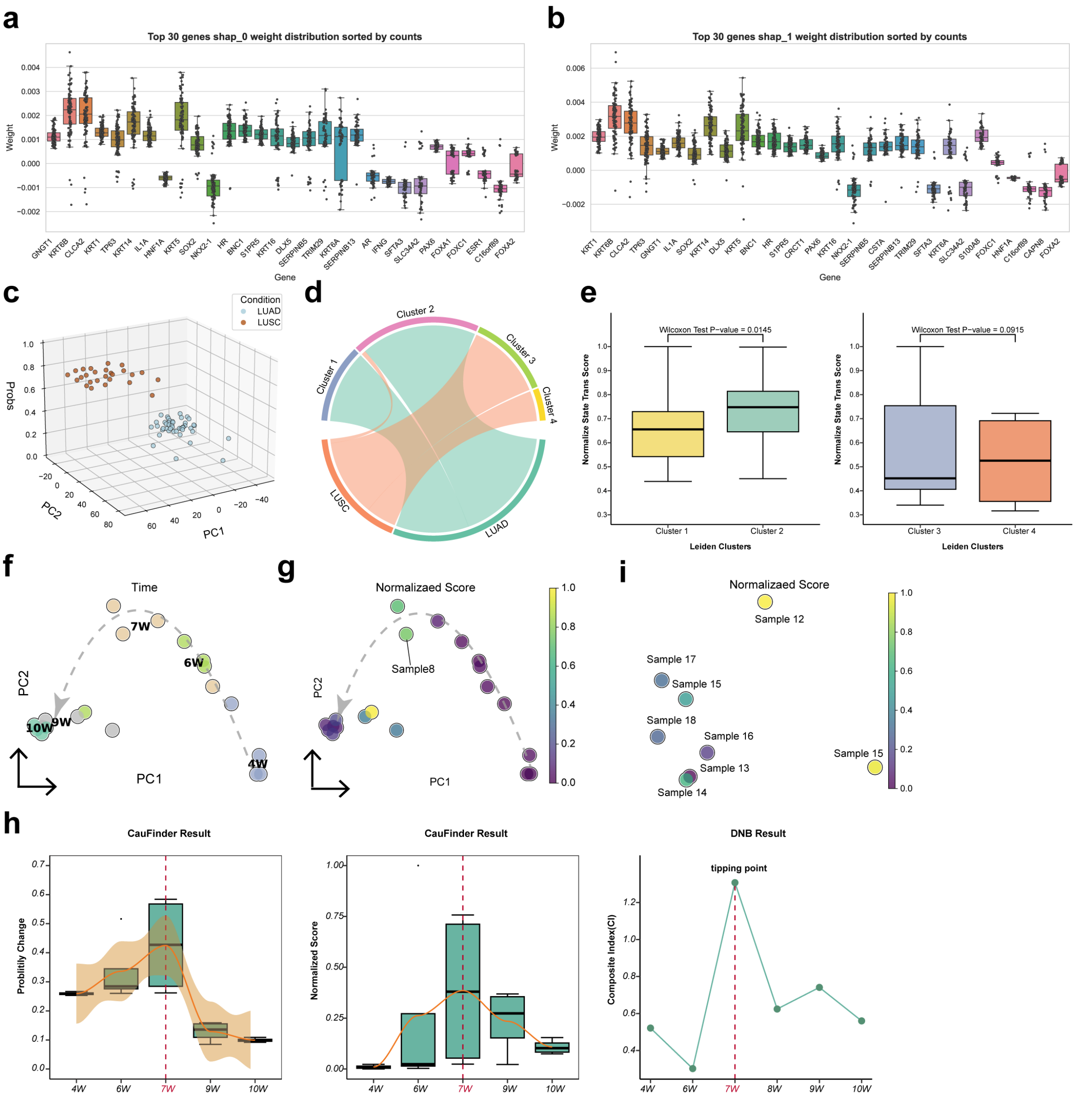

**Supplementary Fig. 9: Identifying causal drivers igniting the transdifferentiation between lung adenocarcinoma and squamous cell carcinoma. a-b**, Boxplots with individual data points showing the distribution of causal weight scores for the top 30 genes. These genes were selected based on their frequency of occurrence across 100 runs. The horizontal axis lists the genes, and the vertical axis represents the causal weight scores derived from the CauFinder model. (**a**) Transition from lung adenocarcinoma (LUAD) to lung squamous cell carcinoma (LUSC). (**b**) Reverse transition from LUSC to LUAD. **c**, 3D visualization of all LUAD and LUSC samples using principal component analysis (PCA). The x and y axes represent PC1 and PC2, respectively, while the z-axis shows the state score of each sample. **d**, Chord plot shows the clustering of samples using the Leiden algorithm with a resolution of 0.25. **e**, Boxplot shows the normalized state transition scores calculated by CauFinder during the simulated state transitions for LUAD samples (Cluster 1 and Cluster 2) and LUSC samples (Cluster 3 and Cluster 4), grouped by cluster. The P value is indicated at the top (based on the Wilcoxon rank-sum test). **f-g**, PCA visualization of RNA-seq data revealing the trajectory of phenotypic transitions from LUAD to LUSC. Each node represents one sample, with the time points (weeks) of sampling shown (**f**), and state transition score of samples on the same PCA embedding (**g**). **h**, Boxplots showing model-derived state probability transitions (left panel) and controllability scores (middle panel) at different time points. The line graph (right panel) displays the composite index (CI) for quantifying the tipping point of system state, based on previous research. The 7-week mark (7W) is highlighted as the tipping point. **i**, State transition scores of samples at 9 and 10 weeks on the PCA embedding, illustrating the transition from LUSC to LUAD.

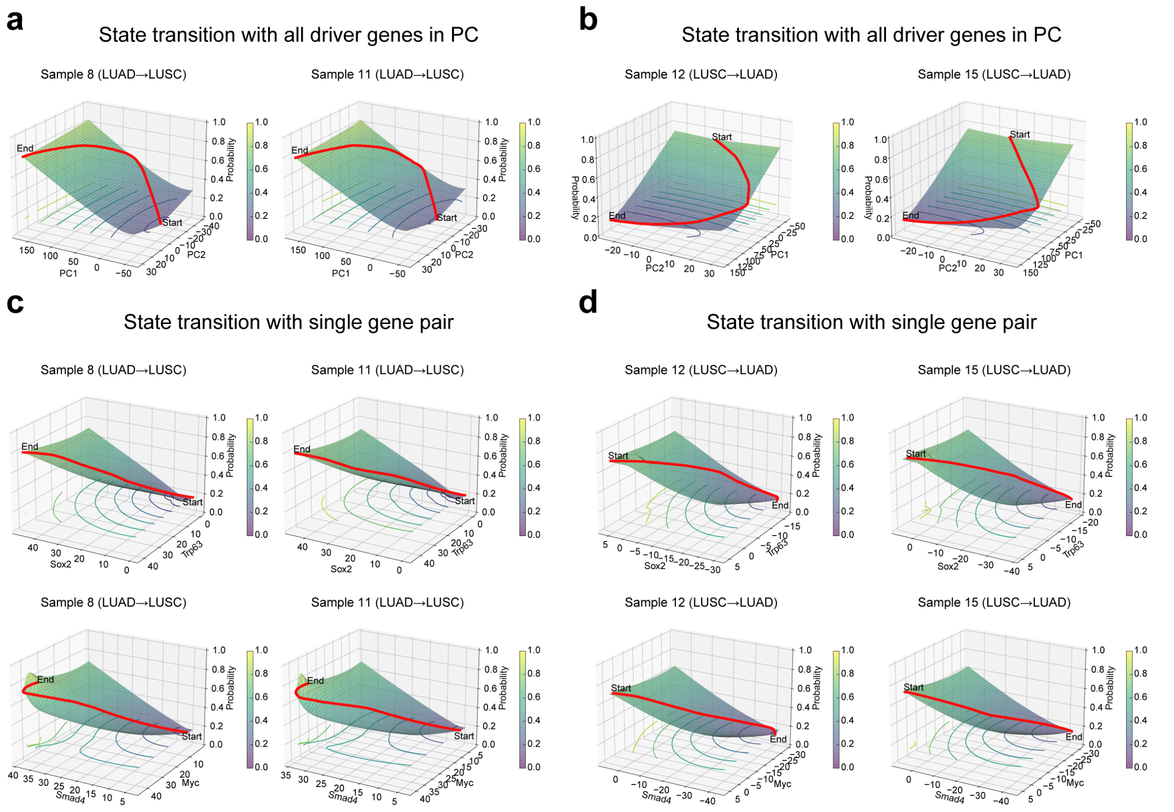

**Supplementary Fig. 10: Computational simulation of the state transitions between LUAD and LUSC, controlled by drivers identified through CauFinder.** The figure is organized to showcase both the natural transition from LUAD to LUSC (**a**, **c**) and the reversed transition from LUSC to LUAD (**b**, **d**). Panels **a** and **b** represent the transitions controlled by all drivers identified by CauFinder after PCA dimensionality reduction, offering a comprehensive view of how these drivers influence the state changes. In contrast, panels **c** and **d** focus on transitions governed by specific gene pairs. The upper panels (**c** and **d**) highlight gene pairs that have been experimentally validated in previous studies, while the lower panels showcase our newly identified gene pair, Myc and Smad4. Each panel plots gene expression against a state probability score as defined by CauFinder, where values approaching 0 indicate proximity to LUAD, and values nearing 1 denote proximity to LUSC.

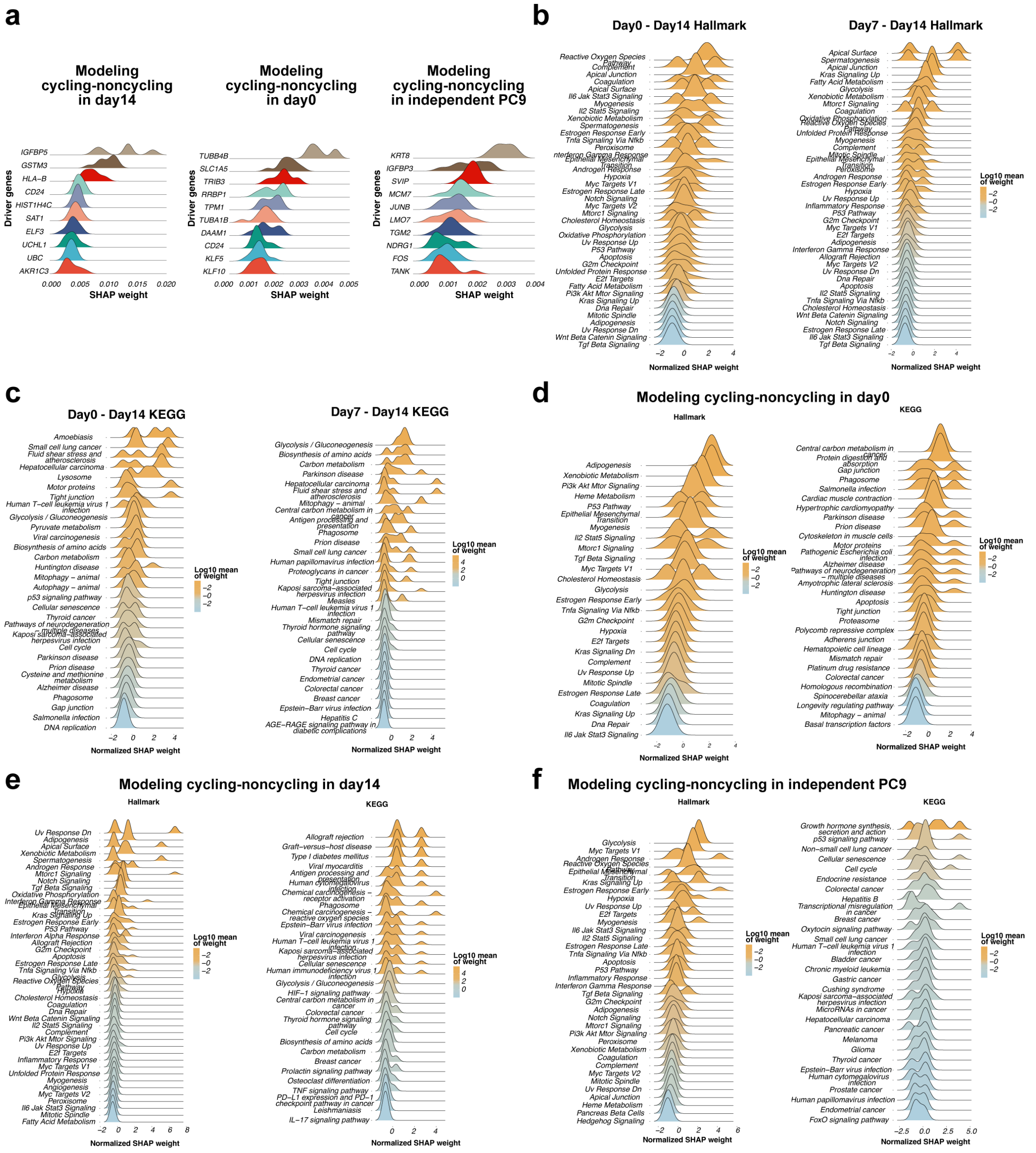

**Supplementary Fig. 11: Causal genes identified by CauFinder and enrichment analysis results in cycling persister cells arising. a,** The ridge plot shows the weights of the top 10 drivers predicted by CauFinder across ten calculations for the groups: day 14 (left), day 0 (middle), and EGFR-driven lung cancer (PC9) as an independent supplement (right). Persister cycling cells and non-cycling counterparts are labeled as 1 and 0, respectively. **b-c,** Weight distribution of all causal drivers included in the HALLMARK pathways (**b**) and KEGG pathways (**c**) in day0 -day14 paired input CauFinder model (left) and day7-day14 paired input CauFinder model (right), ranked by the median weight of causal drivers within each pathway. All weights are log-transformed. **d-f,** Weight distribution of all causal drivers included in the KEGG pathways and HALLMARK pathways in three different cycling-noncycling pairs: day0 (**d**), day14 (**e**), additional PC9 cell line (**f**).

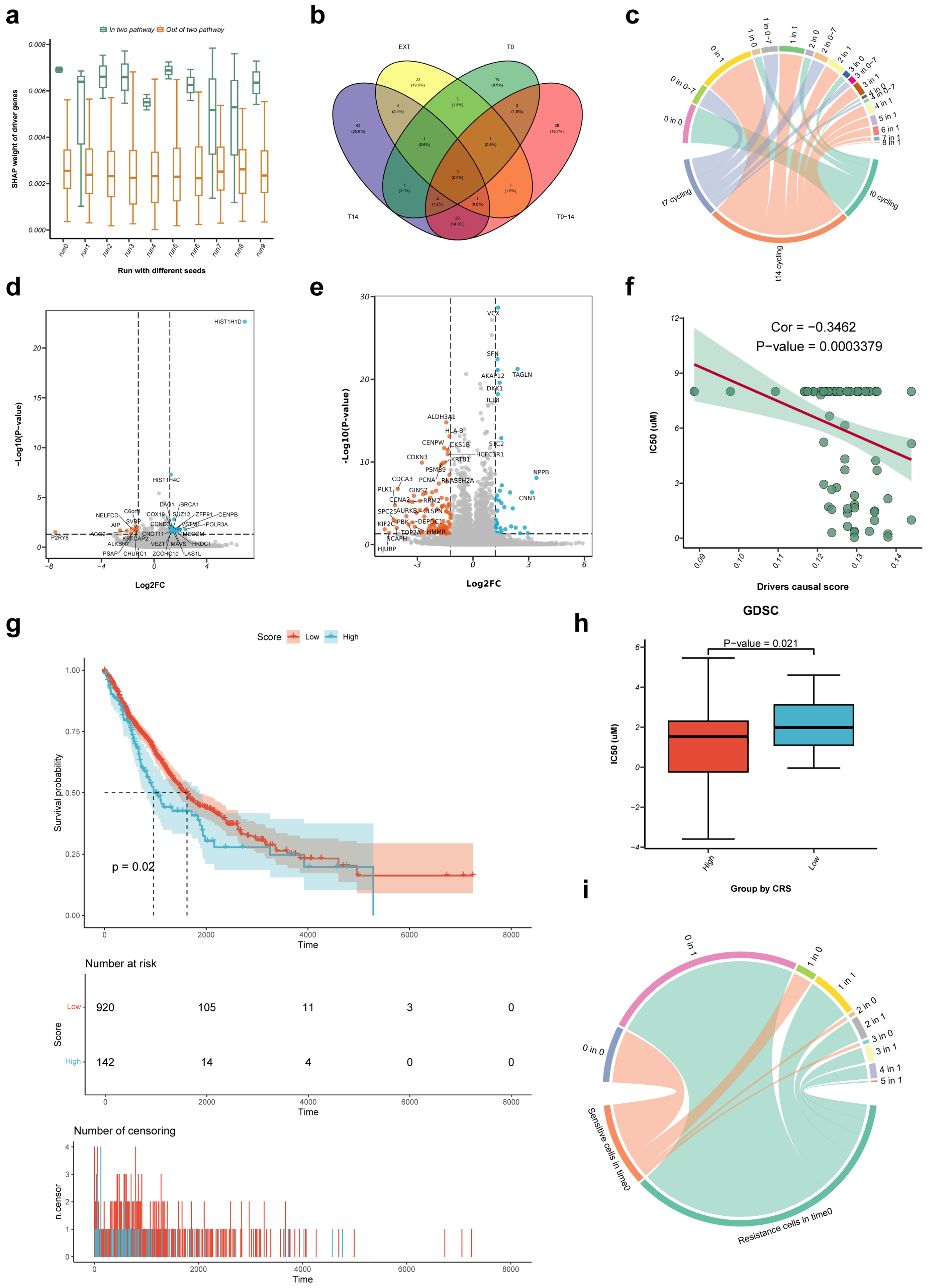

**Supplementary Fig. 12: Downstream analysis on single-cell datasets based on causal genes identified by CauFinder. a,** Box plots showing the weights of two groups of causal drivers across ten runs. The causal drivers were identified using ten different random seeds with fixed CauFinder inputs (day0-day14 paired) and parameters. The drivers were grouped based on their presence in the ROS and FAM pathways. **b,** The Venn plot illustrates the intersection of all drivers predicted by CauFinder, using pairs labeled as 1 for persister cycling cells and 0 for non-cycling counterparts as input. **c,** The chord plot shows the correspondence between persister cycling cells with different time points sampling and the merged pesudo-cells. The pesudo-cells which used as input for the state transition simulation by CauFinder, are displayed at the top, while the original persister cycling cells which categorized by sampling time are shown at the bottom. **d,** Volcano plot showing differential expression genes in pseudo-cell with highest state transition score and other pseudo-cells in persister cycling cells which sampling in day0. **e,** Volcano plot showing differential expression genes in pseudo-cell with lowest state transition score and other pseudo-cells in persister cycling cells which sampling in day14. Blue dots represent up-regulated genes in HST-pCell and red dots down-regulated genes. **f,** Correlation between causal driver scores (x axis) and IC50 (y axis) in patients (dots) from CCLE. **g,** The causal driver scores were associated with the survival probability of lung cancer patients (N = 1062). A log rank test was used to assess the significance between two curves, with a p-value =0.02. **h,** Boxplots showing the distribution of IC50 of Osimertinib treated patients in GDSC across two groups categorized by the median causal driver scores. **i,** The chord plot shows the correspondence between cells with different drug response and the merged pesudo-cells. The pesudo-cells which used as input for the state transition simulation by CauFinder are displayed at the top, while the original cells which categorized by drugs response are shown at the bottom.

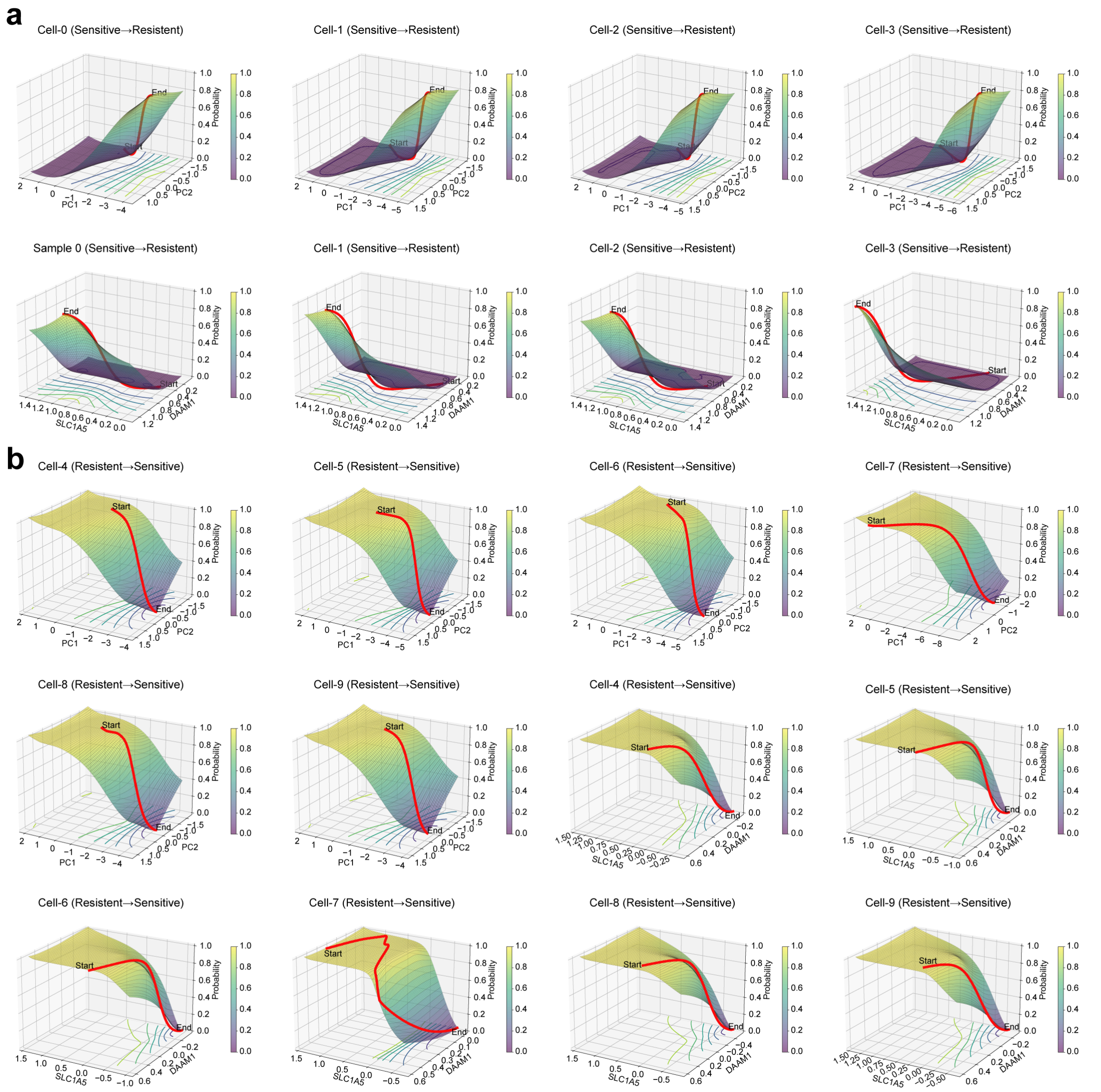

**Supplementary Fig. 13: Computational simulation of state transition via control of causal genes in pseudo-cells.** **a,** Simulation of state transition from drug-sensitive to resistant cells, using either all causal drivers (shown by PCA) or a combination of single biologically significant features (DAAM1 and SLC1A5). The upper panel shows the transition trajectory using PCA, while the lower panel focuses on the specific features. **b**, Simulation of reverse state transition from drug-resistant to sensitive cells, using the same methods as in panel (**a**). The upper panel uses PCA, and the lower panel highlights the single significant features.

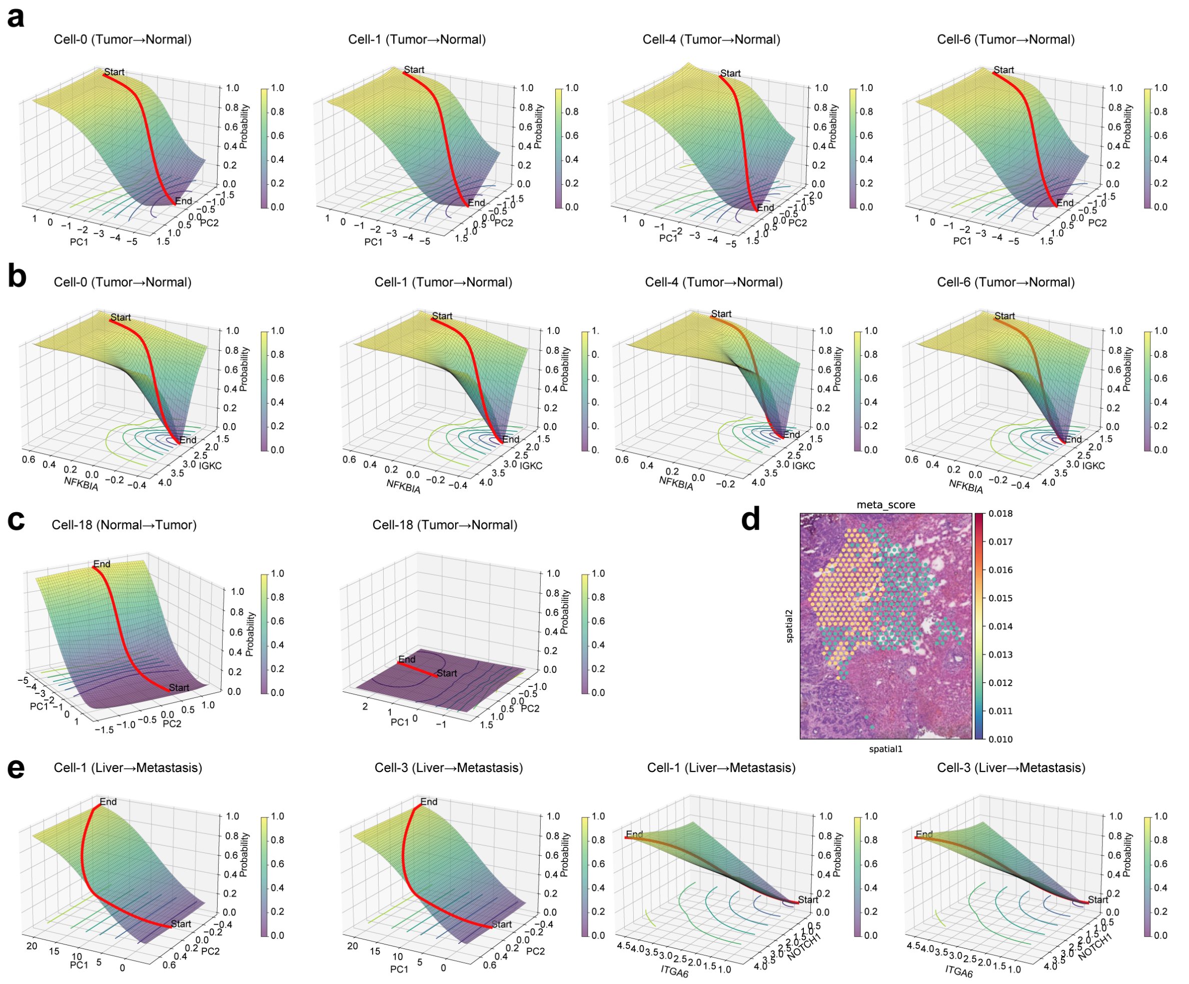

**Supplementary Fig. 14: Computational simulation of state transition of pseudo-cells between tumor and adjacent normal tissues, and between metastatic cancer and nearby liver tissues. a-b,** Computational simulation of decreasing state transition from tumor to normal via control of causal genes for pseudo-cells from P1 tissue, based on either all drivers (shown by PCA, **a**) or a combination of single biologically significant features (NFKBIA and IGKC, **b**). **c,** Computational simulation of decreasing state transition from tumor to normal via control of causal genes for cell-18 based on all drivers (shown by PCA). **d,** Scatter plot in spatial coordinates showing the state transition score based on pseudo-cells at the tumor-liver tissue interface. **e,** Computational simulation of increasing state transition via other pseudo-cells not mentioned in **Fig. 5o**.

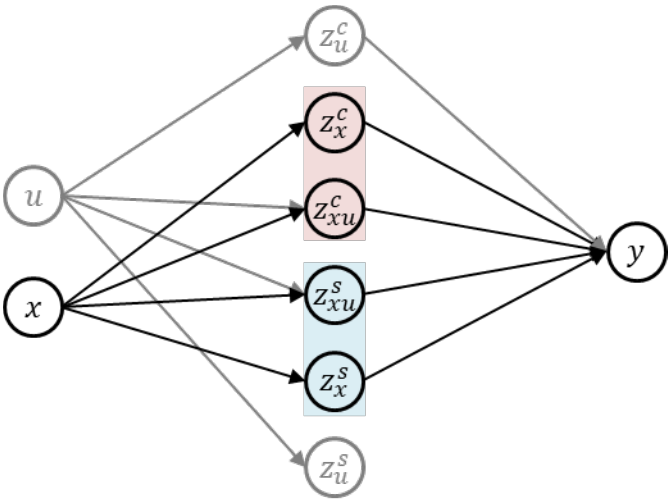

$$P(y|do(z^c)) = \sum_{x,u} P(y|z^c, x, u)P(x, u)$$

**Supplementary Fig. 15: Structural causal model with unobserved variables.** In the causal graph, $x$ represents observed variables, and $u$ represents unobserved variables, which are further decomposed into causal and spurious components within the latent space. The unified causal latent spaces, $z^c = \{z_{xu}^c,\ z_x^c\}$, consist of $z_{xu}^c$, influenced by both $x$ and $u$, and $z_x^c$, influenced solely by $x$. Similarly, the unified spurious latent spaces, $z^s = \{z_{xu}^s,\ z_x^s\}$, include $z_{xu}^s$, affected by both $x$ and $u$, and $z_x^s$, affected only by$x$. The complete set of latent variables, $z = \{z^c,\ z^s,\ z_u^c,\ z_u^s\}$, encompasses all components derived from $x$ and $u$.
